# Human milk oligosaccharide metabolism by *Clostridium* species suppresses inflammation and pathogen growth

**DOI:** 10.1101/2025.01.21.633585

**Authors:** Jonathan A. Chapman, Andrea C. Masi, Lauren C. Beck, Hannah Watson, Gregory R. Young, Hilary P. Browne, Yan Shao, Raymond Kiu, Andrew Nelson, Jennifer A. Doyle, Pawel Palmowski, Márton Lengyel, James P. R. Connolly, Christopher A. Lamb, Andrew Porter, Trevor D. Lawley, Lindsay J. Hall, Nicholas D. Embleton, John D. Perry, Janet E. Berrington, Christopher J. Stewart

## Abstract

Gut microbiome development is strongly impacted by breastmilk and human milk oligosaccharides (HMOs), with short- and long-term health implications. HMO metabolism is best characterised within bifidobacteria, but the full range of other HMO-utilising species remains unknown. This study examined HMO-utilising bacteria that colonise preterm infants (born <32 weeks’ gestation), their role in microbiome modulation, and their effects on intestinal barrier function. We found that infant-derived *Clostridium perfringens*, and three other *Clostridium* species, metabolise HMOs, producing short chain fatty acids, tryptophan catabolites, and other metabolites known to improve host health. *C. perfringens* inhibited pathobiont growth and supressed inflammation in preterm-derived intestinal organoids. These findings suggest a previously unrecognised role for *C. perfringens*, specifically those lacking the perfringolysin O gene, in promoting healthy gut development during infancy.

## Introduction

Early life gut microbiome development plays a critical role in shaping short- and long-term health. Preterm infants born <32 weeks of gestation undergo altered development of their gut microbiome that is linked to pathologies such as necrotising enterocolitis (NEC), an inflammatory mediated bowel disease with a high risk of mortality and morbidity(*1*). Receipt of human milk is an important driver of infant gut microbiome composition(*2*) and protects against NEC, most likely through provision of bioactive factors, such as human milk oligosaccharides (HMOs)(*3–7*). HMOs are complex unconjugated sugars indigestible to humans that act as prebiotics for some gut bacteria, most notably *Bifidobacterium* spp. that are associated with breast-fed babies’ microbiome and health(*8*).

HMO utilising bacteria such as *Bifidobacterium* spp. improve intestinal barrier function and positively influence immune system development, lowering systemic inflammation(*9*), and protecting against immune-mediated diseases such as atopy and asthma(*10*). This is partly mediated by bacterial metabolites including short-chain fatty acids (SCFAs)(*11*) and tryptophan catabolites(*12, 13*) that interact with host cells. *Bifidobacterium* spp. also shape the wider microbiota by producing metabolic breakdown products from HMOs, allowing “cross-feeding” by other beneficial species, promoting their growth and suppressing growth of pathogens(*14, 15*). This knowledge has contributed to the rise in probiotic use in preterm infants over the last decade and inclusion of synthetic HMOs in term formula, including 2′-Fucosyllactose (2′-FL), Lacto-N-tetraose (LNT), lacto-N-neotetraose (LNnT), and 6′-Sialyllactose (6′SL)(*16*). Recent studies have demonstrated other genera can also digest HMOs through varied pathways, as observed for *Bacteroides*(*17*)*, Akkermansia*(*18*) and *Roseburia-Eubacterium* group(*19*). Notably, these genera are rare in preterm infants(*20*) and the full diversity of bacteria that metabolise HMOs within this population is unknown.

The current study sought to comprehensively describe novel HMO-utilising bacteria that colonise preterm infants, their role in modulating the microbiome, and their impact on intestinal barrier function as proposed mechanisms of neonatal health.

## Results

### *Clostridium spp.* and *Bifidobacterium* spp. isolated from preterm infant stool can metabolise HMOs

We screened the abilities of 29 bacterial isolates, mostly from preterm infant stool, to grow on six different HMOs, and glucose and lactose (Fig. 1A). These species were obtained by untargeted cultivation and represent a median of 80% (interquartile range 61%-91%) of all relative microbial abundance observed in preterm infants(*20*). Only *Bifidobacterium* and *Clostridium* species were able to use HMOs (Fig. 1A). Except for *B. animalis*, all *Bifidobacterium* (n = 7 isolates) grew on at least one of the HMOs, with LNT and LNnT being most frequently used. Except for *C. butyricum,* all *Clostridium* species, namely *C. perfringens, C. tertium*, *C. baratii* and *C. paraputrificum* (n = 11 isolates), were able to grow on one or more of the HMOs tested.

**Figure 1.**
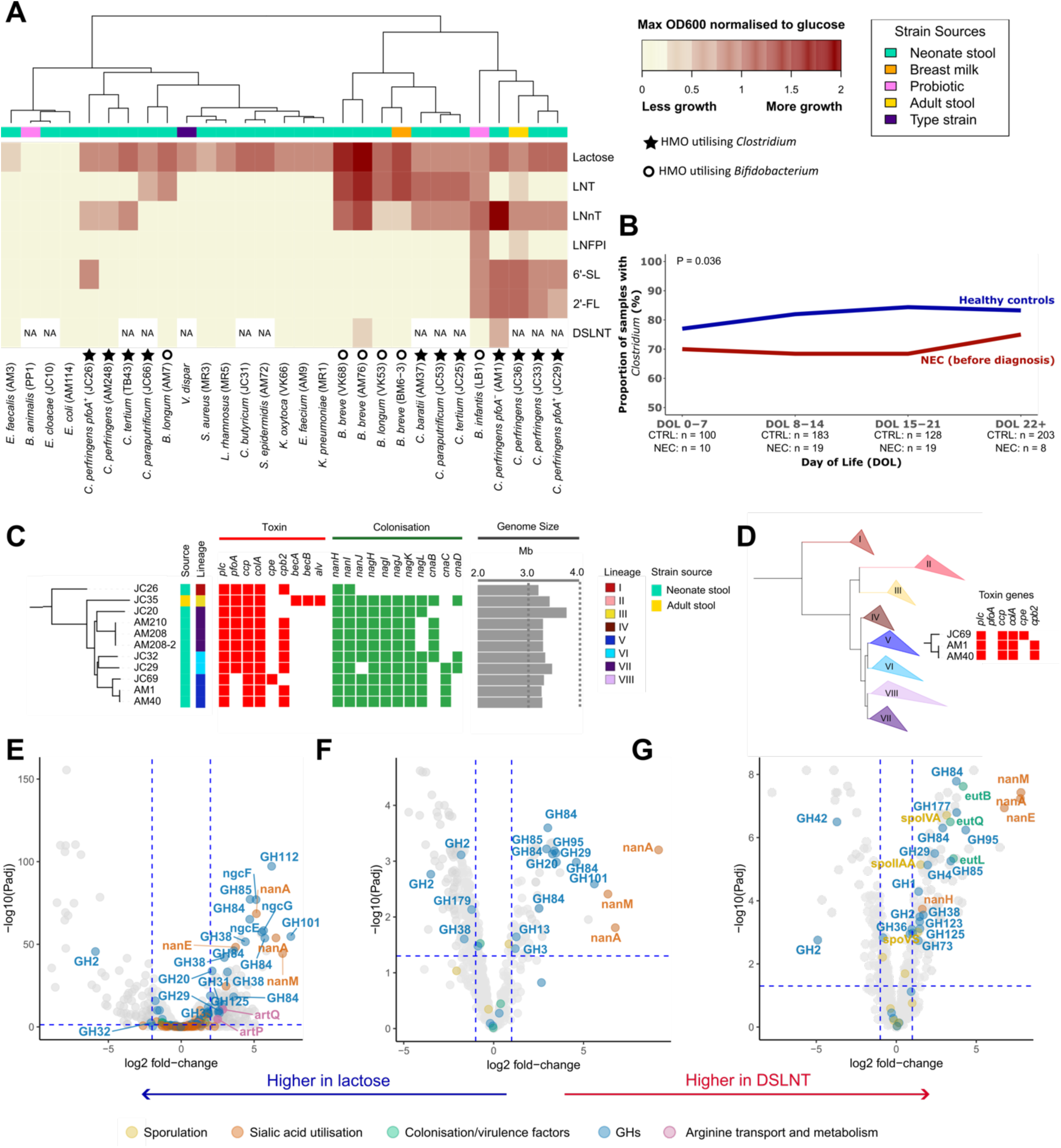
*Clostridium* spp. and *Bifidobacterium* spp. have the capacity to metabolise human milk oligosaccharides (HMOs). (**A**) Heatmap representing the growth of 29 bacterial isolates on six HMOs and lactose. The values reported represent the maximum OD600 reached normalised to glucose. (**B**) *Clostridium* prevalence in infants who developed NEC (n= 22) and control (n= 179) infants from the publicly available data from the MAGPIE study(*22*). Control infants had significantly higher prevalence of *Clostridium* compared to samples from NEC infants before diagnosis (P=0.036). (**C**) Alignment of 11 novel genomes with toxin and colonisation profiles and genome sizes. Lineage assignment is according to Kiu *et al*. (2023)(*21*). (**D**) A phylogenetic tree of 313 strains where the 11 novel genomes in this study were compared with 302 strains assigned to 8 lineages, as in Kiu *et al*. (2023)(*21*). Lineage V is a hypovirulent clade. Three isolate genomes in this study were assigned to this lineage. All three were found to be lacking toxin gene *pfoA*, which is typical of lineage V strains. (**E-G**) Volcano plot of RNA-srq data (**E**) and proteomics data on supernatant (**F**) and pellet (**G**) for AM1 grown on DSNT vs lactose. A positive log2 fold-change indicates upregulation on DSLNT relative to lactose, while a negative fold change indicates downregulation on DSLNT relative to lactose. HMO, human milk oligosaccharide; LNT, lacto-N-tetraose; LNnT, lacto-n-neotetraose; LNFP I, lacto-N-fucopentaose I; 6’-SL, 6’-sialyllactose; 2’-fucosyllactose,; DSLNT, disialyllacto-N-tetraose; NEC, necrotising enterocolitis; GH, glycoside hydrolase.

Four *C. perfringens* isolates clustered based on sugar growth profile with the probiotic derived *B. infantis*. Having discovered *C. perfringens* could use HMOs we tested a wider collection of *C. perfringens* strains (obtained from Kiu *et al.,* 2023(*21*)) which showed all could use HMOs (fig. S1). Unlike *Bifidobacterium*, *C. perfringens* generally could not grow on LNT. However, compared to all other isolates, *C. perfringens* strain AM1 showed the best growth on disialyllacto-N-tetraose (DSLNT), an HMO consistently associated with protection from NEC (*5, 7*) (Fig. 1A).

Using genus level publicly available data from the MAGPIE study(*22*), we found *Clostridium* was prevalent in preterm infants, but at significantly lower rates in those diagnosed with NEC compared with time-matched healthy controls (p = 0.036; Fig. 1B). At the strain level, *C. perfringens* lacking the gene encoding toxin perfringolysin O (*pfoA*) are linked to the neonatal-derived health-associated hypovirulent lineage V(*21*), while carriage of *pfoA* is associated with increased risk of NEC and paediatric inflammatory bowel disease(*23*). Thus, in subsequent work we focused on AM1, a *C. perfringens pfoA* negative isolate notable for its ability to grow on DSLNT (this specific AM1 isolate will be referred to as CP-*pfoA^−^* henceforth) (Fig. 1A, 1C, and 1D). This isolate was also found to be susceptible to commonly prescribed antibiotics in neonatal intensive care (table S1).

To identify genes potentially involved in HMO metabolism, RNA-Seq was performed during exponential growth of CP-*pfoA^−^* using DSLNT, LNnT, 6’SL and lactose. Transcriptome data showed clustering based on sugar growth profiles (fig. S2A). Comparing each HMO to lactose, the highest number of differentially expressed genes (DEGs) were observed with DSLNT, followed by LNnT, and 6′SL (table S2). Among the top 20 upregulated DEGs with DSLNT were those encoding a predicted glycoside hydrolase (GH) 101 cazyme (endo-α-N-acetylgalactosaminidase; locus 01633) which showed the highest log fold change (7.5), and enzymes involved in sialic acid metabolism (*nanM*, two *nanA* genes) (Fig. 1E). Specific to DSLNT, the most upregulated genes included one encoding a GH112 protein (1,3-beta-galactosyl-N-acetylhexosamine phosphorylase, locus 02923), and three genes involved in diacetylchitobiose transport (*ngcG*, *ngcF*, *ngcE*; loci 02918, 02919, 02920). On 6′SL, the top 20 upregulated DEGs involved sialic acid (*nanM*, two *nanA*, *nanE*) and fucose (*fucI*, *fucU*, *fucA*, *fucO*, *fucP*) metabolism (fig. S2B). Finally, in LNnT, most top upregulated genes encoded hypothetical proteins, but also genes involved in arginine metabolism and transport (*argG*, *argH*, *artQ*, *artP*) (fig. S2C). A total of 11 different classes GHs were upregulated across all HMOs tested, three of which have been shown to act on HMOs; GH29, GH85, and GH112 (*24*). Others, like the GH84 *nagJ* have been shown to act on mucin O-glycans and may also target HMOs owing to structural similarity(*25*).

Proteomics on supernatant (i.e., secreted) and cell pellet (i.e., intracellular or cell-associated) peptides also clustered by sugar utilised (fig. S2E). Similar to RNA-Seq, growth on DSLNT resulted in upregulation of multiple GH family and sialic acid degrading (NanA, NanM, NanE, NanH) proteins (Fig. 1F). The protein encoded in locus 01633 (top upregulated gene in transcriptomics) was the 4th most significant protein in DSLNT supernatant, but not in the pellet, suggesting that this is an uncharacterised enzyme acting extracellularly, likely in conjunction with two NanA proteins and NanM which represented the three most upregulated proteins. HMO quantification of spent media showed that lactose, LNnT, and 6′SL were completely degraded, but DSLNT metabolism generated undigested byproducts, in particular LNT (table S3). Specifically, accumulation of LNB suggests the presence of an enzyme able to break the β 1-3 bond between LNB and lactose prior to complete desialylation. The presence of LNT further indicates that the sialidases could act on both DSLNT and the sialylated LNB, and the full digestion of DSLNT was possible only when the β 1-3 bond was cleaved first.

### *Clostridium* spp. produced wider varieties and higher quantities of beneficial metabolites than *Bifidobacterium* spp

We next compared metabolites produced by HMO-utilising *Clostridium* (*C. perfringens*, *C. tertium*, *C. baratii* and *C. paraputrificum*) and *Bifidobacterium* (*B. infantis*, *B. breve* and *B. longum*) strains. We hypothesised that *Clostridium* species metabolising HMOs may, like *Bifidobacterium* species, produce beneficial metabolites such as SCFAs that play critical roles in gut health including providing energy, regulating the immune system, maintaining gut barrier integrity, and modulating microbiome composition(*11, 26*). We therefore quantified SCFAs within culture supernatants of strains grown with individual HMOs, glucose or lactose. Compared with *Bifidobacterium,* supernatants from *C*. *perfringens* and *C. baratii* contained significantly greater growth adjusted concentrations of butyrate (all p <0.05), and propionate (all p <0.05) (Fig. 2A). *C. perfringens* and *C. baratii* also produced greater total quantities of SCFAs than *Bifidobacterium* (Fig. 2A & 2B). Consistent findings were seen in media supplemented with either glucose or lactose (fig. S3).

**Figure 2.**
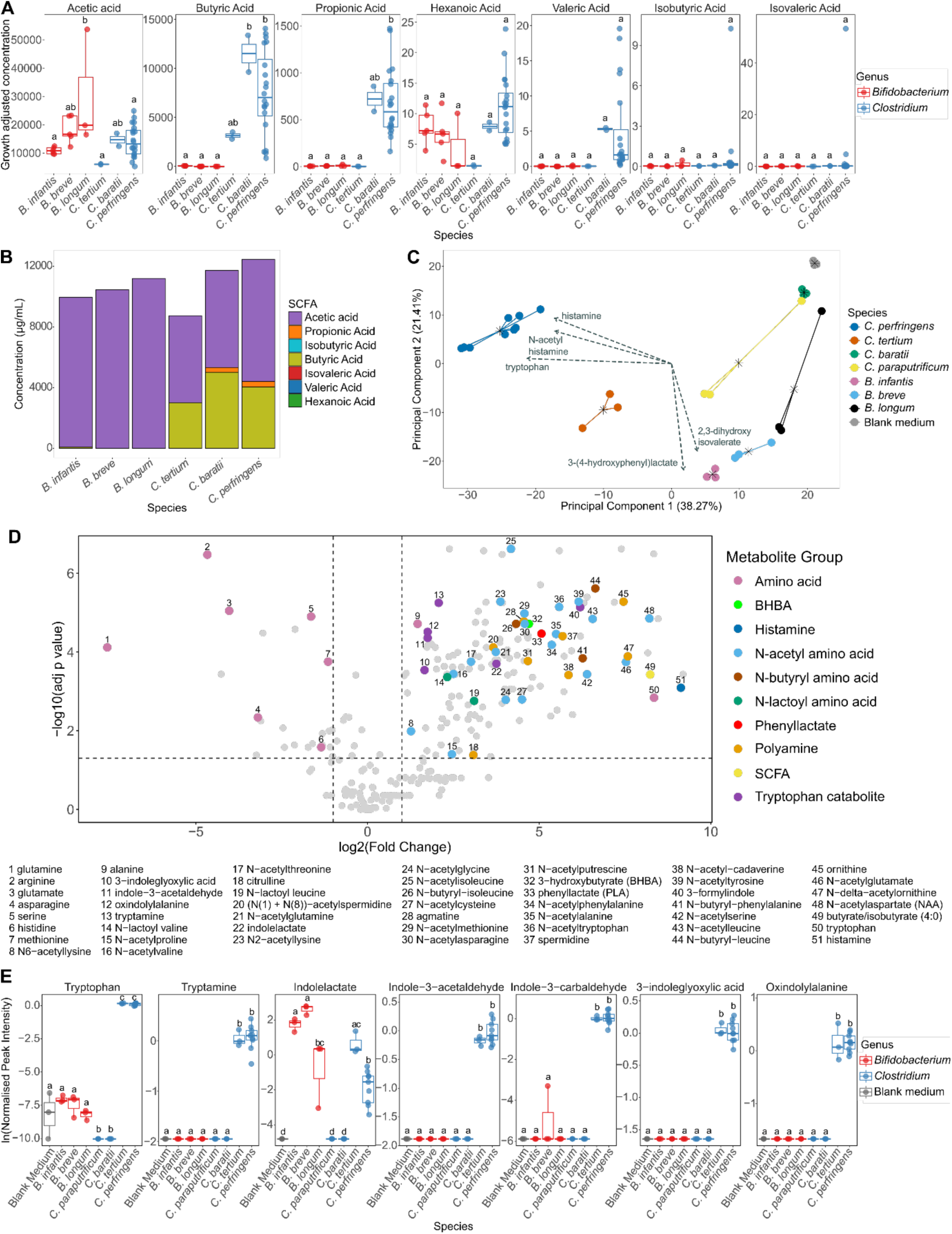
*Clostridium* spp. produced wider varieties and higher quantities of beneficial metabolites compared to *Bifidobacterium* spp. (**A**) Box plots showing individual short chain fatty acid (SCFA) production by species grown on HMOs. SCFA concentrations in μg/mL were divided by the maximum OD600 recorded for each strain to provide growth adjusted concentrations. Conditions with the same letters are not significantly different. (**B**) Stacked bar plots showing the total concentration of SCFAs in culture supernatants of *Bifidobacterium* and *Clostridium* spp. grown on individual HMOs. (**C**) Principal component analysis (PCA) of untargeted metabolomics data for cell free supernatants (CFSs) generated from bacterial cultures growing on cocktails of HMOs. Arrows indicate the top five metabolites by loadings magnitude. (**D**) Volcano plot comparing the metabolites detected in CFSs of *C. perfringens pfoA^−^* (AM1) growing on a cocktail of the HMOs 6’-SL, LNnT and 2’-FL, compared with blank ZMB1 medium. A positive log2 fold-change indicates production of metabolites by the strain, while a negative fold change indicates metabolite depletion. Metabolites of interest are highlighted and numbered. (**E**) Box plots showing the levels of tryptophan catabolites produced by *Bifidobacterium* and *Clostridium* spp. during growth on HMOs. Conditions with the same letters are not significantly different. SCFA, short chain fatty acids; BHBA, beta-hydroxybutyric acid.

To investigate a broader range of metabolite production, we next generated cell free supernatants (CFSs) from each strain of interest and performed untargeted metabolomics. Two CFSs were produced per strain, using either a strain-specific mixture of utilisable HMOs or glucose. Unsupervised ordinations of the metabolomic profiles following growth on HMOs showed clustering by species and more broadly by genus (Fig. 2C). This was consistent with glucose (fig. S4A), and glucose and HMO metabolomic profiles were comparable within each strain (table S4). Thus, we focused subsequent analysis on CFSs derived from growth with HMOs.

Compared to media controls, CP-*pfoA^−^* CFS showed significantly increased polyamine formation, tryptophan biosynthesis and catabolism, butyrate/isobutyrate production, and generation of other neuromodulatory, immunomodulatory, or antimicrobial metabolites, with depletion and modification of amino acids (Fig. 2D). Many of these metabolites were differentially abundant in other CFSs, with high similarities between the three *C. perfringens* and *C. tertium* CFSs (fig. S4C-J). *B. infantis* (LB1; Labinic™, Biofloratech Ltd) and *B. breve* (AM76) CFSs were distinct in containing multiple gamma-glutamyl amino acids (fig. S4F-G). *C. baratii* (AM37), *C. paraputrificum* (JC53) and *B. longum* (AM7) showed a low number of these metabolites of interest, namely five, three and zero respectively (fig. S4H-J).

*C. perfringens* and *C. tertium* produced a broad range of indole-containing tryptophan catabolites (associated with promoting intestinal barrier function and inhibiting inflammation)(*12, 27, 28*) along with tryptophan itself, at significantly higher levels than *Bifidobacterium* and other *Clostridium* spp. (Fig. 2E). Additionally, *C. perfringens* and *C. tertium* CFSs contained higher levels of polyamines(*29*) (associated with increased tight junction expression and inflammation reduction)(*30, 31*) and their precursors (fig. S5A). Only ornithine and citrulline were significantly raised in *Bifidobacterium* CFSs, specifically in *B. infantis* (LB1). Only *C. perfringens* CFSs contained significantly raised levels of histamine (produced by probiotic strains, can alter intestinal motility(*32*), and suppress cytokine secretion and wider intestinal inflammation(*33*)) (fig. S5B). Neuromodulators 3-hydroxybutyrate and N-acetylaspartate were also significantly higher in *C. perfringens* CFSs compared to all other strains (fig. S5B). Finally, phenyllactate (broad spectrum antimicrobial) was significantly higher in all *Bifidobacterium* and *Clostridium* CFSs compared to media only, except for *C. baratii* (fig. S5B).

### *Clostridium* spp. CFS suppressed the growth of pathobionts and promoted the growth of naturally occurring *Bifidobacterium* spp

We next assessed whether the *Clostridium* and *Bifidobacterium* CFSs could suppress the growth of four of the most abundant pathobionts present in the preterm gut microbiome (*Escherichia coli*, *Klebsiella pneumoniae*, *Klebsiella oxytoca* and *Enterobacter cloacae*)(*20*), all of which were isolated from preterm infant stool (Fig. 3). Each CFS showed inhibitory activity against at least three pathobionts, with the majority suppressing growth of all four (Fig. 3A-B). CFS from the probiotic *B. infantis* had the strongest inhibitory activity, with growth of all pathobionts reduced to <2% of their no treatment controls. *C. tertium* and three *C. perfringens* CFSs showed a similar pattern of high inhibition and clustered with *B. infantis* based on CFS inhibitory capacity (Fig. 3A). These four *Clostridium* CFSs reduced each pathobiont’s growth to ≤55% of their no treatment controls (all p <0.001). *K. pneumoniae* was most susceptible to the four *Clostridium* and *B. infantis* CFSs, with growth reduced by each to <15% of the no treatment control (all p <0.001).

**Figure 3.**
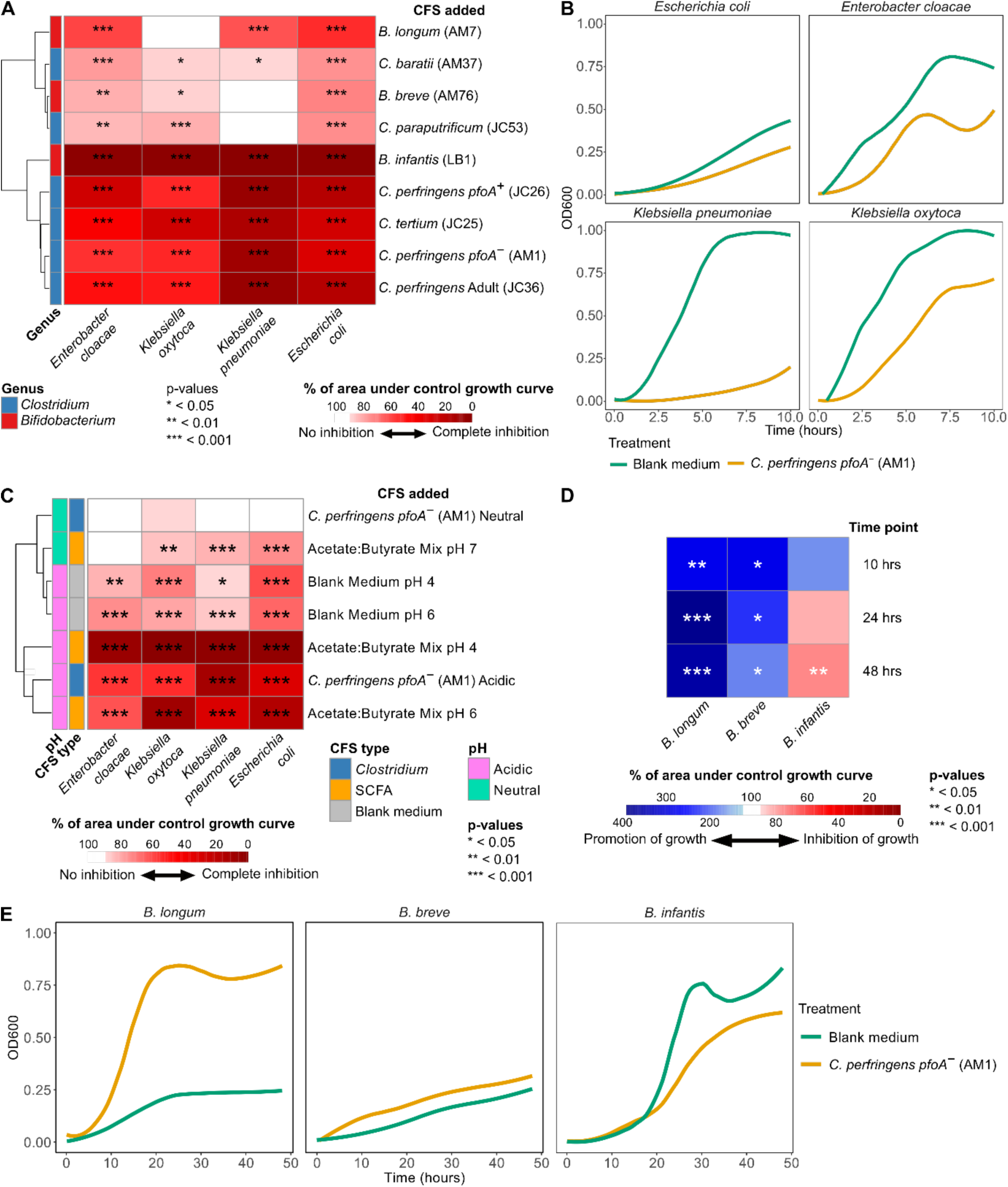
*Clostridium* spp. cell free supernatants (CFSs) suppressed pathogen growth without impacting naturally occurring *Bifidobacterium* spp. Growth. (**A**) Growth of pathobionts in ZMB1 supplemented with glucose with CFSs from six *Clostridium* and three *Bifidobacterium* isolates. Values represent area under the curve (AUC) for growth in media supplemented with CFS as a percentage of the control AUC. (**B**) Growth curves of pathobionts in ZMB1 supplemented with glucose and treated with AM1 CFS. (**C**) Impact of acidic pH and/or the presence of the SCFAs acetate and butyrate on pathobiont growth. Values represent AUC for growth as a percentage of the control AUC. (**D**) Growth of *Bifidobacterium* isolates in ZMB1 supplemented with glucose and AM1 CFS. Values represent AUC for growth in media supplemented with CFS as a percentage of the control AUC. (**E**) Growth curves of *Bifidobacterium* spp. in ZMB1 supplemented with glucose and treated with AM1 CFS.

All CFSs were found to be weakly acidic, with mean pH ranging from 6.64 to 4.36 (fig. S6A). Additionally, all strains used to generate CFSs are prolific producers of acetate and, in the case of the *Clostridium*, also butyrate (Fig. 2C). We therefore speculated that pH or SCFAs may be mediating the inhibitory effects seen. Pathobiont growth was tested with an acetate:butyrate mix, blank ZMB1 medium and CP-*pfoA^−^* CFS, all adjusted to multiple pHs. Acidic blank medium inhibited growth, but not to the same extent as CFSs, while acidic SCFAs matched or exceeded the inhibitory activity of CFSs (Fig. 3C). Neutralised SCFAs also inhibited three pathobionts, but to a lesser extent, while neutralised CP-*pfoA^−^* CFS lost its activity. The most pH sensitive of the pathobionts, *E. coli* (Fig. 3C), was then used to test the effect of adjusting pH on all CFSs. Only three CFSs, including CP-*pfoA^−^*, lost their activity when neutralised, while the remainder had reduced activity (fig. S6B). The loss of inhibition following CP-*pfoA^−^* CFS neutralisation suggests pH dependence, likely mediated by pH sensitive metabolites, such as SCFAs. The similar reduction in inhibitory effects observed with neutralised SCFA mix and CP-*pfoA^−^* CFS further supports this role. However, acidic blank medium was less inhibitory than CP-*pfoA^−^* CFS so pH is not the sole factor and additional pH-sensitive antimicrobial factors likely also contribute to the inhibitory effect.

Within the neonatal gut, it is critical that beneficial bacteria are not inhibited during suppression of pathobiont growth. The CFS of CP-*pfoA^−^* was selected to determine its impact on *Bifidobacterium* species growth owing to it being in the hypovirulent linage V, lacking the *pfoA* toxin gene, its ability to use health associated DSLNT (*5–7*) and its production of several beneficial SCFAs and metabolites. CP-*pfoA^−^* CFS derived from growth on glucose was found to significantly enhance the growth of naturally occurring *B. breve* and *B. longum* across all time points (all p<0.05; Fig. 3D-E). However, a significant impact on the growth of the probiotic (i.e., not naturally occurring) *B. infantis* was found after 48 hours (p<0.01; Fig. 3D-E). Notably, because the CP-*pfoA^−^* CFS used in these experiments was derived following growth on glucose, the promotion of *B. breve* and *B. longum* is due to microbial metabolites and not cross-feeding of HMO degradation byproducts.

### *Clostridium* CFSs dampened inflammation in an intestinal organoid model

Diet-microbe-host interaction in the preterm gut is critical in understanding health and disease and developing effective therapies(*34*). We examined the potential toxicity of the CFSs harvested following growth on HMOs on the human gut using Caco-2 cells before exposing preterm infant-derived intestinal organoids (PIOs). When added to a concentration of 25% v/v, the majority of *Clostridium* spp. and *Bifidobacterium* spp. derived CFSs did not reduce Caco-2 viability below 50% (fig. S7). This concentration was therefore selected for subsequent experiments. Whilst there was strain-to-strain variability, *Clostridium* CFSs were less toxic than *Bifidobacterium* CFSs at 25% and 50% v/v. Notably, CP-*pfoA^−^* CFS at 10% and 25% v/v maintained full Caco-2 viability (fig. S7).

The preterm gut epithelium was modelled using PIO monolayers, within an anaerobic co-culture system(*35*) (Fig. 4A). PIOs were treated with selected CFSs in isolation and in the presence of inflammatory stimuli (LPS and flagellin). CP-*pfoA^−^* CFS was tested alongside CFS from hypervirulent linage I *C. perfringens* JC26 (CP-*pfoA^+^*), *C. tertium* (related HMO-using *Clostridium* spp.), and *B. infantis* (commercially available probiotic strain).

**Figure 4.**
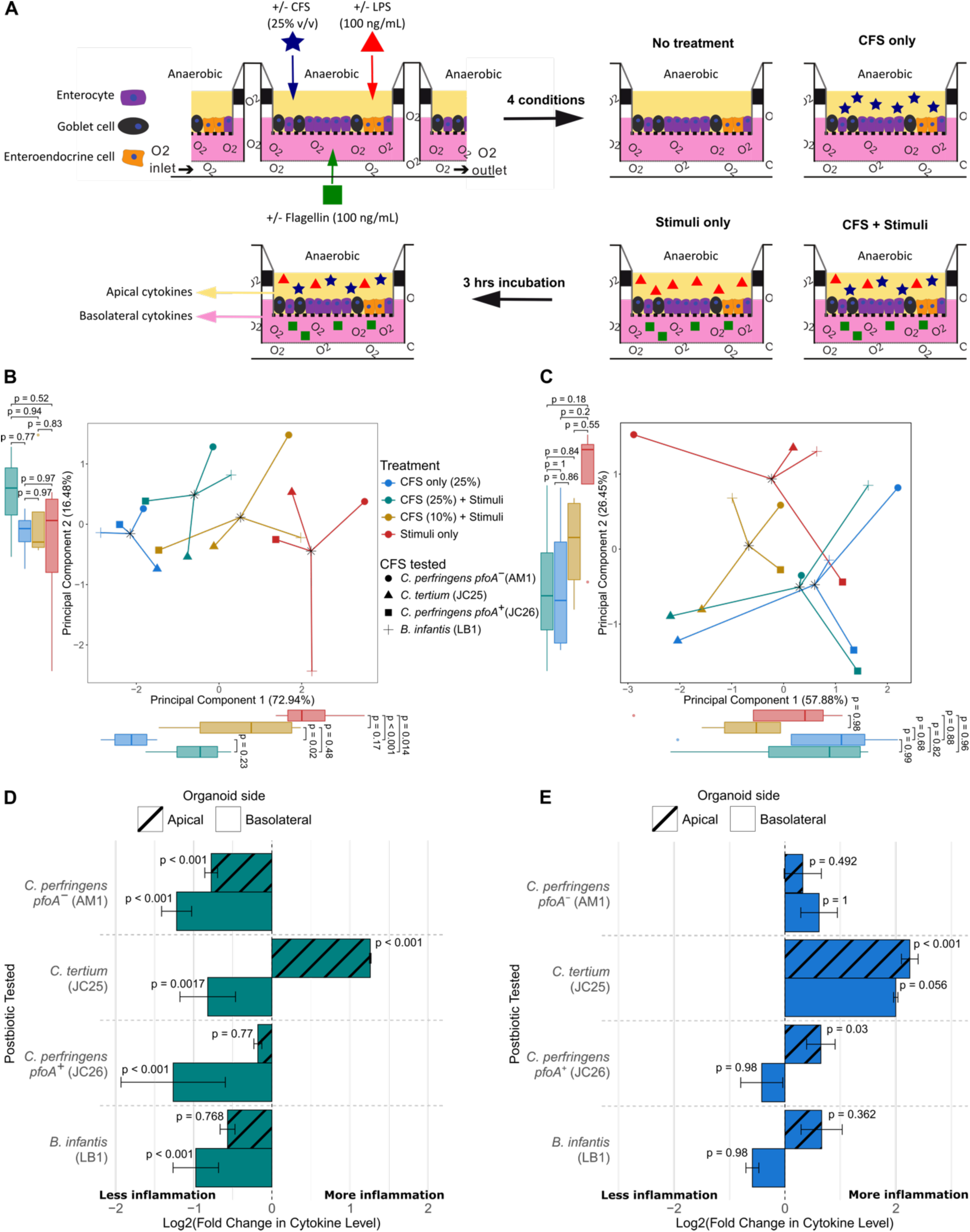
*Clostridium* spp. and *Bifidobacterium* spp. cell free supernatant (CFS) dampened inflammation in a preterm intestinal-derived organoid co-culture model. (**A**) Schematic of organoid monolayer inflammation assays. (**B-C**) Principal component analysis (PCA) of basolateral (B) and apical (C) cytokine (IL8, TNF-κ, CCL2, CCL7 and CXCL5) profiles. The data were converted to fold changes compared to the negative control and transformed. There was no apical CXCL5 data for the experiment with the AM1 cell free supernatant (CFS) so this cytokine was removed from apical analysis. (**D**) Bar plots showing log2 fold change of apical and basolateral IL8 secretion during combined “CFS + Stimuli” compared to the “Stimuli only”. P values represent the differences between the unprocessed detected cytokine levels. (**E**) Bar plots showing log2 fold change of apical and basolateral IL8 secretion during “CFS only” compared to the “No treatment”. P values represent the differences between the unprocessed detected cytokine levels.

PIOs lack immune cells but secrete cytokines that can be used as a proxy for immunomodulation. Unsupervised ordination of basolateral cytokine data showed overall cytokine profiles of PIOs treated with LPS and flagellin (‘Stimuli only’) were significantly different from those treated with CFS only regardless of the specific strain (all principal component 1 (PC1) p <0.05; Fig. 4B). In the presence of stimuli, only treatment with 25% v/v CFS resulted in cytokine profiles that were significantly different from ‘Stimuli only’ (PC1 p = 0.014) and comparable to ‘CFS only’ controls (PC1 p = 0.23; Fig. 4B). This trend was not observed in the apical cytokine data (Fig. 4C).

All CFSs significantly inhibited stimuli-induced basolateral secretion of IL8 (all p <0.05; Fig. 4D; full datasets in tables S5-8). Only CP-*pfoA^−^* also significantly inhibited IL8 apically (p <0.001), while *C. tertium* induced significantly greater apical secretion (p <0.001). Changes in secretions of a further four inflammatory cytokines were also detected (fig. S8A-D). Only CP-*pfoA^−^* and CP-*pfoA^+^* inhibited basolateral secretion of all four, with only CP-*pfoA^−^* also significantly inhibiting apical secretion of both CCL7 (p <0.001; fig. S8B) and CCL2 (p <0.001; fig. S8D). *C. tertium* (p <0.001) and CP-*pfoA^+^* (p = 0.02) induced significant increases in apical TNF-α. We confirmed that the inhibitory effects observed were not caused by the ZMB1 medium used to produce the CFSs, except in the case of apical CCL2 (p = 0.035; fig. S9; table S9). Notwithstanding, the fold change was lower than with CP-*pfoA^+^*, *B. infantis*, and CP-*pfoA^−^*, indicating components of ZMB1 media were not solely responsible. Except for basolateral IL8, where inhibition was reduced at neutral pH, inhibition patterns were comparable between acidic and neutral pH CP-*pfoA^−^* CFSs, confirming that the dampening of pro-inflammatory cytokines was not due to the acidic pH of the CFSs (fig. S10; table S10).

We also tested whether CFSs at 25% v/v alone could induce inflammatory cytokine secretion compared to PIO medium only. Both CP-*pfoA^−^* and *B. infantis* CFSs did not induce any significant IL8 secretion, but CP-*pfoA^+^* (p = 0.03) and *C. tertium* (p <0.001) both triggered apical IL8 production (Fig. 4E). For the other four cytokines measured, *C. tertium* induced apical CXCL5 (p = 0.026), TNF-α (p <0.001), and CCL7 (p <0.001), CP-*pfoA^+^* induced apical TNF-α (p = 0.017), *B. infantis* induced apical CCL7 (p = 0.04), while CP-*pfoA^−^* did not induce secretion of any cytokine apically or basolaterally (fig. S11A-D). Thus, overall only CP-*pfoA^−^* CFS had no proinflammatory impact.

### The presence of *pfoA* is critical in determining the impact of live *C. perfringens* and its CFS on the intestinal epithelium

Aside from CFS, we next assessed the impact of live CP-*pfoA^−^* combined with inflammatory stimuli on PIOs. Apical secretion of CCL2, CCL7 and CXCL5 and basolateral secretion of IL8 were all significantly reduced in the presence of live CP-*pfoA^−^* (all p ≤ 0.005; Fig. 5A; table S11), indicating similar activity to the CFS. CP-*pfoA*^+^ and CP-*pfoA*^−^ share four toxin and two colonisation genes, with CP-*pfoA*^−^ encoding a further seven colonisation factors (Fig. 1C). Proteomics confirmed CP-*pfoA*^+^ CFS contained PfoA, while it was absent from CP-*pfoA*^−^ (Fig. 5B). Toxicity assays in Caco-2 cells showed CP-*pfoA*^+^ CFS reduced cell viability to 49%, compared with 99% for CP-*pfoA*^−^ (p <0.001; Fig. 5C). In PIO monolayers under aerobic conditions, CP-*pfoA*^+^ CFS increased non-mitochondrial oxygen consumption (p <0.001), reduced ATP production (p = 0.054) and increased proton leak (p <0.001) (Fig. 5D, fig. S12). CP-*pfoA*^−^ CFS also increased proton leak to a similar degree (p <0.001), but simultaneously enhanced basal respiration (p <0.001), maximal respiration (p <0.001), ATP production (p = 0.012) and spare respiratory capacity (p = 0.002). Overall, this analysis revealed CP-*pfoA*^+^ CFS and CP-*pfoA*^−^ CFS reduced and increased mitochondrial bioenergetic function, respectively.

**Figure 5.**
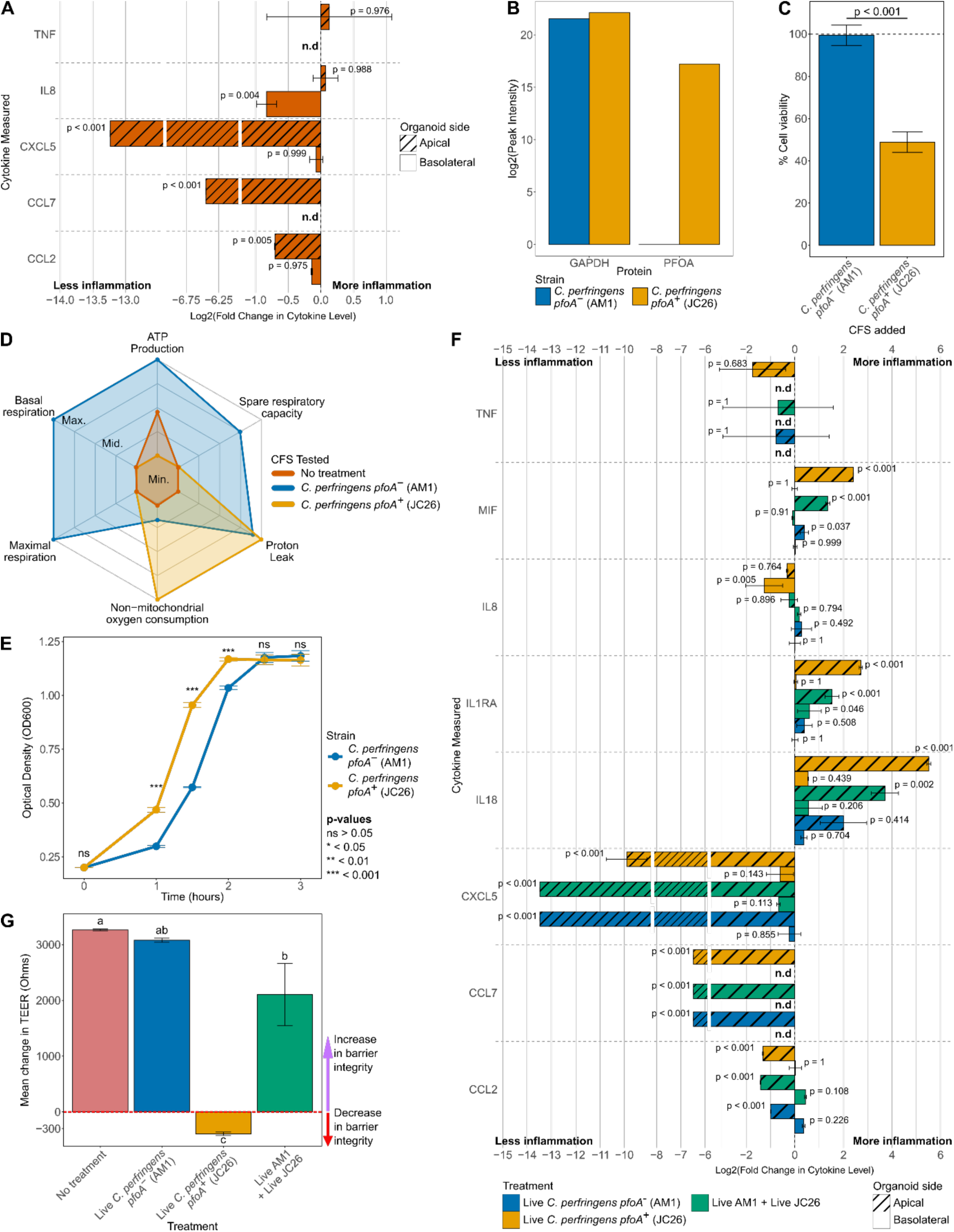
*Clostridium perfringens* showed strain specific impacts on preterm intestinal-derived organoid monolayers during live co-culture depending upon *pfoA* carriage. (**A**) Bar plots showing log2 fold change in apical and basolateral cytokine secretion during combined treatment with live *C. perfringens pfoA^−^* (AM1) for 3 hrs, with inflammatory stimuli added at 1 hr, compared to “Stimuli only”. P values represent the differences between the unprocessed detected cytokine levels. (**B**) Proteomic detection of PFOA in the *C. perfringens pfoA^−^* (AM1) and *C. perfringens pfoA^+^* (JC26) cell free supernatant (CFS) used in co-culture experiments, with GAPDH provided for reference. (**C**) MTS tetrazolium dye cell viability assay data for Caco-2 cells treated with 25% v/v CFSs from *C. perfringens pfoA*^−^ (AM1) and *C. perfringens pfoA^+^* (JC26) strains. Values are shown as % of the viability measured for Caco-2 cells incubated in regular DMEM. (**D**) Radar plots showing the summary of changes to mitochondrial energetic function induced by CFSs, quantified as changes to oxygen consumption rate (OCR) of cells. Detailed quantification of changes to each parameter and associated statistical analyses are shown in fig. S12. (**E**) Monoculture growth curves of *C. perfringens pfoA^−^* (AM1) and *C. perfringens pfoA^+^* (JC26) growing in ZMB1 medium supplemented with glucose. (**F**) Changes in apical and basolateral cytokine secretion during treatment with either live *C. perfringens pfoA^−^* (AM1), live *C. perfringens pfoA^+^* (JC26), or live *C. perfringens pfoA^−^* (AM1) + *C. perfringens pfoA^+^* (JC26). Data has been converted to fold change compared to the “No treatment” condition and log2 transformed for plotting. P values represent the differences between the unprocessed detected cytokine levels. (**G**) Change in barrier integrity as measured by trans-epithelial electrical resistance (TEER) during organoid incubation with live *C. perfringens* strains. Conditions with the same letters are not significantly different. CFS, cell free supernatant; TEER, trans-epithelial electrical resistance; n.d, not detected

We next sought to compare microbe-microbe and microbe-host interaction with live co-culture of CP-*pfoA*^+^ and CP-*pfoA*^−^ on PIOs. We observed CP-*pfoA*^+^ had a faster growth rate than CP-*pfoA*^−^ (Fig. 5E). Cytokine secretion from PIOs treated with live CP-*pfoA*^−^ and CP-*pfoA*^+^ in the anaerobic co-culture system were then assessed against untreated controls. Live *C. perfringens* significantly reduced apical CCL7, CXCL5, and CCL2 secretion to below basal levels, regardless of *pfoA* status (all p ≤ 0.001; Fig. 5F, table S11). Live CP-*pfoA^+^* halved basolateral IL8 (p = 0.005) and more than quadrupled apical MIF, IL1RA, and IL18 (all p <0.001). CP-*pfoA*^−^ also increased MIF (p = 0.037), but to lower levels than CP-*pfoA*^+^, and had no significant impact on any other cytokine compared to basal media.

Given the contrasting results between potentially beneficial CP-*pfoA*^−^ and pathogenic CP-*pfoA^+^*, we next investigated whether pre-colonisation of PIOs with CP-*pfoA*^−^ could protect against CP-*pfoA^+^* damage. This pre-colonisation with CP-*pfoA*^−^ significantly reduced the levels of MIF, IL1RA, and IL18 that are elevated by CP-*pfoA*^+^ (all p ≤ 0.002; fig. S13A-C). Finally, live CP-*pfoA*^+^ also significantly compromised barrier integrity (p <0.001), which was prevented by pre-colonisation with CP-*pfoA*^−^ (p = 0.003; Fig. 5G). In isolation, CP-*pfoA^−^* increased barrier integrity, with transepithelial electrical resistance (TEER) values similar to untreated PIOs (p = 0.906).

## Discussion

Using preterm stool bacterial isolates, we discovered numerous *Clostridium* species from the preterm infant gut microbiome can metabolise HMOs*. C. perfringens* showed strain-to-strain variation in HMO use independent of *pfoA* status, but only a *pfoA^−^* strain could use DSLNT, a HMO that is linked to reduced NEC risk(*5–7*). Compared to *Bifidobacterium*, *Clostridium* spp. produced a broader range of SCFAs including butyrate, as well as a greater diversity of tryptophan catabolites and other potentially beneficial metabolites. CP-*pfoA^−^* CFS showed inhibitory effects on common infant pathobionts, generally promoted growth of beneficial microbes, enhanced mitochondrial bioenergetic function in preterm intestine-derived organoids, and suppressed the inflammatory response in an organoid co-culture model. When using live microbes to study competitive exclusion, CP-*pfoA^−^* protected against CP-*pfoA^+^* mediated damage to epithelium barrier integrity and suppressed the pro-inflammatory activity of the pathogenic strain. Thus, CP-*pfoA^−^* may play an important and previously unrecognised role in gut microbiome mediated immune education during early life.

*C. perfringens* are especially prevalent in newborns(*21, 36*), which may explain their ability to utilise HMOs. Given evolution has favoured HMOs to be highly abundant in human milk, early life colonisers that use these prebiotics have been proposed as likely having therapeutic potential, as seen with probiotics containing HMO using *Bifidobacterium* spp.(*37–39*). Indeed, we reported higher prevalence of *Clostridium* in healthy preterm infants from a recent UK-wide study. In contrast, some studies have linked the relative abundance of *Clostridium* to NEC(*40, 41*). Notably, these studies were relatively small (i.e., 8-11 NEC cases) and relied on amplicon sequencing that is limited to the genus level. In recent work focusing on culturing *C. perfringens* strains, *pfoA^+^ C. perfringens* caused cellular damage and the hypovirulent lineage V, notable for the absence of *pfoA,* did not encode the necessary virulence traits required to cause NEC(*21*). Similar findings have been reported in other inflammatory conditions including paediatric inflammatory bowel disease(*23*). Taken together, this underscores the need for strain-level microbiome data and highlights the beneficial potential of CP-*pfoA*^−^.

While no consistent agent has been associated with NEC in preterm infants, *Klebsiella* has been implicated in several studies, including the largest current analysis of NEC infant stool that employed metagenomics(*42, 43*). *C. perfringens, C. tertium,* and *B. infantis* CFSs showed inhibition of *K. oxytoca* and *K. pneumoniae*, as well as other pathobionts. In the case of CP-*pfoA^−^*, this was dependent on acidic pH and likely mediated by SCFAs, although other mechanisms, such as production of bacteriocins, may also be involved. CP-*pfoA^−^* further promoted the growth of naturally occurring infant *B. breve* and *B. longum*, but not commercial probiotic derived *B. infantis* of unknown origin. In industrialised nations *B. breve* and *B. longum* are the predominant *Bifidobacterium* in infants(*2, 20, 44*) and were notable for their strong growth on LNT. Cross-feeding with *C. perfringens,* that metabolised DSLNT into LNT, could therefore be an important contributor to promoting *B. breve* and *B. longum* colonisation in the infant gut.

Probiotics appear to reduce NEC in preterm infants, although precise mechanisms remain unclear(*45*). Competitive exclusion, whereby two species cannot coexist if they have identical niches, may be one mechanism. We found prior colonisation of PIOs with CP-*pfoA^−^* can protect the intestinal epithelium from CP-*pfoA^+^* mediated damage, suggesting CP-*pfoA^−^* could have probiotic potential. However, live biotherapeutic products may carry risks(*46*) and alternative non-viable microbial-derived therapies (e.g., postbiotics) may hold promise. For this reason and considering the recent FDA guidance against using probiotics in preterm infants, we focused on CFS. CFS from CP-*pfoA^−^* had positive roles in modulating the gut microbiome (inhibiting pathobionts and promoting *Bifidobacterium*), improving mitochondrial function (SRC and respiration rate), and suppressing inflammation. This therapeutic potential warrants further exploration for novel microbial therapy based on *pfoA^−^ C. perfringens.* On the other hand, we corroborate and extend previous work through showing that human intestinal cells treated with PfoA-containing CFS from CP-*pfoA^+^* had a ∼50% reduction in viability and showed signs of mitochondrial damage(*21, 47*). Such evidence cautions against supplementing HMOs to infants colonised with *pfoA^+^ C. perfringens*, which is especially important as several commercial formula products now include selected synthetic HMOs in term formula.

In summary, we report that a range of *Clostridium* species use HMOs in the preterm gut, producing a broad range of SCFAs, tryptophan catabolites, and other potentially beneficial immunomodulatory metabolites that influence host physiology. However, where infants are potentially colonised with pathogenic *pfoA^+^ C. perfringens,* these results caution against widespread HMO supplementation. This work further provides a workflow for the discovery of novel microbial therapies and demonstrates the potential of *pfoA^−^ C. perfringens* as a live biotherapy or postbiotic for preterm infants at risk of NEC.

## Materials and methods

### Ethics and sample collection

Preterm infants (born at <32 weeks gestation) were born or transferred to a single tertiary level Neonatal Intensive Care Unit in Newcastle upon Tyne, UK, and participated in the Supporting Enhanced Research in Vulnerable Infants (SERVIS) study (REC10/H0908/39) after written informed parental consent. Stool samples were regularly collected from nappies/diapers of preterm infants into sterile collection pots by nursing staff. Breast milk samples were collected from residuals from an infant’s feeding systems. Samples were initially stored at −20°C before being transferred to −80°C for long term storage. Intestinal tissue samples used to generate organoid cell lines were salvaged following surgical resection.

### Bacterial isolation and identification

Stool samples were thawed on ice and initially diluted roughly 1:10 w/v in sterile anaerobic phosphate buffered saline (PBS). 10-fold serial dilutions were then performed using sterile anaerobic PBS and various dilutions (typically 10^−2^ and 10^−4^) were cultured by adding 100 μl of inoculum onto an agar plate, before spreading to cover the surface of the agar plate and incubating for up to 96 hours. Numerous different agar media were used including brain heart infusion (BHI), Transoligosaccharide propionate (TOS), Bifidus Selective Medium (BSM), De Man, Rogosa and Sharpe (MRS), fastidious anaerobe agar (FAA), and yeast extract-peptone-dextrose (YPD). Unique appearing colonies based on morphology, colour, and size were sub-cultured twice. Full-length 16S rRNA gene sequencing (27F 5′-AGAGTTTGATCCTGGCTCAG3’; 1492R 5′-GGTTACCTTGTTACGACTT-3′) and matrix-assisted laser desorption ionization-time of flight mass spectrometry (Bruker MALDI-TOF MS) of single fresh colonies were used to initially identify isolates to genus or species level. Isolates were added to glycerol for long term storage at −80°C.

### Whole genome sequencing and genomic analysis

Glycerol stocks of 11 *Clostridium perfringens* strains were streaked out on BHI agar plates and incubated in an anaerobic chamber at 37°C overnight. Single colonies were then picked from these plates, transferred to 5 ml BHI broth and incubated, with shaking at 110 rpm, in the anaerobic chamber at 37°C for 48 hours. Cultures were then centrifuged at 5000 G for 10 minutes at 4°C and the supernatants removed. Genomic DNA was extracted using the MasterPure Complete DNA and RNA Purification Kit (Lucigen). Cell pellets were resuspended in 500 μl PBS, centrifuged at 5000 G for 5 minutes and the supernatant removed. Pellets were then resuspended in 300 μl proteinase K master mix, made up following manufacturer’s protocol using stocks from the kit, and incubated in a heat block at 65°C for 15 minutes, with 10 seconds of vortexing every 5 minutes. The incubation then continued for a further 45 minutes. Tubes were cooled to room temperature and 2 μl RNase A from the kit was added to each. Samples were incubated at 37°C for one hour and then cooled on ice for 5 minutes. 150 μl of kit MPC protein precipitation reagent was added to each tube, followed by vortexing for 10 seconds and centrifuging at 5000G, at 4°C, for 10 minutes. Supernatants were transferred to fresh tubes and 500 μl 2-propanol added to each, with mixing done by inverting 40 times. Samples were then centrifuged at 5000G, at 4°C, for 10 minutes and the supernatants removed. DNA pellets were then washed by adding and removing 1 ml 70% ethanol twice. Tubes were then left open in a laminar flow hood to allow any residual ethanol to evaporate and the DNA pellets to dry. 40 μl nuclease free water was then added, with each then briefly vortexed and left to resuspend at 4°C overnight. Extracted DNA was then transferred to −20°C for storage.

All 11 *C. perfringens* isolates were sequenced on the NovaSeq 6000 system (2 ×151 bp) at the Wellcome Sanger Institute. Genomes were assembled using SPAdes(*48*) draft assembly genomes were quality-checked using checkm v1.1.3(*49*) and GUNC v1.0.5(*50*) ensuring ≥ 90% completeness and ≤ 5% contamination prior to taxonomic assignment (species-level assignment) via gtdb-tk v2.3.2(*51*). Draft genome assemblies were then annotated using prokka v1.14(*52*), with GFF annotated files being used for input for core gene alignment construction via panaroo v1.2.8(*53*). Core gene alignment generated was used to construct a phylogenetic tree using IQ-TREE v2.0.5(*54*) together with 673 public genomes published previously for lineage assignment purpose(*21*). Tree was visualised using iTOL v6.0(*55*). Toxin genes and colonisation factors of *C. perfringens* isolates were screened computationally using ABRicate v1.0.1 (https://github.com/tseemann/abricatev) via TOXIper sequence database (https://github.com/raymondkiu/TOXIper). Genome sizes were calculated using sequence-stats v1.0 (https://github.com/raymondkiu/sequence-stats). Mash-distance sequence tree comprising solely 11 *C. perfringens* isolate genomes was generated via Mashtree v1.2.0(*56*) with default parameters. Distance tree was mid-point rooted and visualised in iTOL v6.0(*55*).

### Isolate growth curves

All isolates were grown overnight in BHI, with the exception of bifidobacteria which were grown in MRS supplemented with L-cysteine HCl (0.05% w/v). The chemically defined medium Zhang Mills Block 1 (ZMB1) was selected for use for this work, as it can support the high density growth of a wide variety of organisms (e.g., *Clostridium*, *Bifidobacterium, Lactobacillus, Klebsiella, Bacterioides*, etc) and is well suited to downstream analytical applications, such as metabolomics(*57*). ZMB1 without glucose was prepared, and various sugars were tested in 1% w/v concentration: base ZMB1 medium, glucose, lactose, 2’FL, DSLNT, LNT, LNnT, LNFP I, 6’SL. All isolates could not be tested on DSLNT due to limited supply of this HMO. The overnight growth for each isolate was centrifuged at 5000 G for 5 minutes at 4°C, and the pellet was resuspended in the same amount of anaerobic PBS. 20 μl of the growth resuspended in PBS was spiked in 180 μl of media, and the growth was measured in a Cerillo Stratus plate reader for 150 hours. All isolates were tested in triplicate, and wells with media only were included in each plate to check for contamination.

### Analysis of *Clostridium* prevalence in an existing preterm infant microbiota dataset

The previously published Mechanisms Affecting Gut of Preterm Infants (MAGPIE) Study recruited preterm infants from NICUs across the UK and generated microbiota data using 16S rRNA gene sequencing (V4 region)(*22*). In total, 56 samples were available prior to NEC diagnosis along with 614 healthy controls (no NEC diagnosis). Details of microbial DNA extraction, library preparation and sequence data processing are available in the original study manuscript. Feature counts of bacterial OTUs in the original MAGPIE data were merged at the genus level to determine the prevalence of *Clostridium* in preterm infant and one sample per participant was selected across pre-defined weekly timepoints based on day of life (DOL): 0-7; 8-14; 15-21; and 22+. Where multiple samples were available for a single patient at any given timepoint, those from the earlier DOL were selected. We used only samples before NEC diagnosis to avoid bias from NEC treatment (e.g., antibiotics) and, given the median day of NEC onset is day 20, we grouped all samples from week 4 onwards. A two-sided z-test was used to test differences in the proportion of samples where *Clostridium* was present between infants diagnosed with NEC and controls using the “prop.test” function in R(*58*).

### Antibiotic resistance testing

Antimicrobial susceptibility testing was performed using the agar dilution method following the CLSI guidelines. Brucella agar (Oxoid, UK) supplemented with 5% laked sheep blood, hemin and vitamin K was used for the tests. The antimicrobial agents tested included vancomycin (range 0.125 to 4 mg/L), ampicillin (0.016 to 4 mg/L), metronidazole (0.064 to 8 mg/L), meropenem (0.004 to 0.5 mg/L), penicillin (0.008 to 1 mg/L). All antibiotics were purchased from Discovery Fine Chemicals. All isolates were grown on preferred agar for 48 hours at 37°C in an anaerobic chamber before testing. For each isolate 4-5 representative colonies were picked and resuspended to 0.5 McFarland bacterial suspension and 1 µL was inoculated using a multipoint inoculator. Plates were incubated in an anaerobic chamber at 37°C for 48 hours. Minimum inhibitory concentrations (MICs) were identified as the lowest concentration of antimicrobial agent leading to visible inhibition of growth compared to control plate without antibiotics. Control strains with defined MIC concentrations were represented *C. perfringens* NCTC 8237, *B. fragilis* NCTC 9343, *S. aureus* NCTC 12973 and *E. coli* NCTC 12241. Resistance to antibiotics was defined based on the EUCAST clinical breakpoints 2024 for clostridia using the *C. perfringens* breakpoints, while for bifidobacteria the breakpoint values used were taken from the data for “anaerobe, gram positive bacteria” reported in EUCAST clinical breakpoints 2021.

### RNA-sequencing of *C. perfringens* AM1 grown on specific HMOs

*C. perfringens* AM1 was grown in BHI to mid-exponential phase. Bacteria were centrifuged at 5000 G for 5 minutes at 4⁰C and then resuspended to 0.1 starting OD in ZMB1 supplemented with either lactose or the HMOs they could grow on at a 0.5% concentration (w/v) (LNnT, DSLNT, 6’SL). RNA was extracted from 2 ml of culture when it reached mid-exponential phase using the RNeasy Mini Kit (Qiagen) following the manufacturer’s instructions. Bacteria were pelleted by centrifuging 8000 G 1 min, supernatant was saved for further analysis, and the cells were resuspended in 700 μl of RLT solution and 500 μl of 100% ethanol. The subsequent steps were performed as per the protocol instructions and the RNA was eluted in 30µl in RNase-free water. Total RNA was quantified using the Qubit™ RNA High Sensitivity kit (Invitrogen™) and the RNA integrity number (RIN) was determined using the RNA 6000 Nano Kit on the BioAnayzer 2100 (Agilent). Total RNA was diluted to 100 ng and ribosomal RNA depletion was carried out using the Ribo Zero Plus kit (Illumina) as per manufacturer’s instructions. Library preparation was performed on the depleted RNA using the NEBNext® Ultra™ II Directional RNA Library Prep Kit for Illumina (New England Biolabs) and sequenced on the NovaSeq 6000 SP 100 cycle kit (Illumina). High-quality reads were aligned to the AM1 strain genome using Bowtie2 (version 2.4.5)(*59*) using the ‘very-sensitive’ option. Aligned reads were then processed using HTSeq (version 2.0.8)(*60*) to generate gene-level count data. “DESeq2” (version 1.44.0)(*61*) in R (version 4.4.0)(*58*) was used for counts normalisation and differential gene expression comparison between conditions. A gene was considered differentially expressed when absolute log2 fold change value > 2 and adjusted P value < 0.05.

### Proteomics of *C. perfringens* AM1 grown on specific HMOs

Proteomics was performed on cell pellets and supernatants from *C. perfringens* AM1 grown to mid-exponential phase. 500 µl of bacterial culture was centrifuged at 8000 G for 1 minute, the supernatant was saved, and the pellet was washed twice in 1 ml of cold PBS. The second PBS wash was removed, and the pellet and supernatant were stored at −80°C until they were analysed. Secretome samples were first precipitated using methanol/chloroform method. To 250 µl of growth media, 528 µl of Methanol and 66 µl of chloroform were added, then vortexed and mixed with 698 µl of water. Samples were vortexed again and centrifuged for 15 minutes at 4000 G, 4°C. The top phase of the supernatant was removed. 1ml of cold methanol was added followed by 30 minutes centrifugation at 16000 G, 4°C. The supernatant was discarded, the pellet air dried and dissolved in 30 µl of S-trap lysis buffer (5%SDS, 50mM TEAB, pH 8.5). The cell pellets were sonicated in 100 µl of S-trap lysis buffer.

Protein concentrations were measured using Micro BCA™ Protein Assay.

Equivalent of 15 µg of total protein was used for digestion. Proteins were reduced with dithiothreitol (DTT) at the final concentration of 20 mM (65⁰C, 30 minutes). Cysteines were alkylated by incubation with iodoacetamide (40 mM final concentration, 30 minutes, room temperature in dark) and then acidified by adding 27.5% Phosphoric acid to a final concentration of 2.5% (v/v). The samples were then loaded onto spin columns in six volumes of binding buffer (90% methanol 100mM TEAB pH 8) and centrifuged at 4000 G for 30 seconds. The columns were then washed with binding buffer (three times) and the flow through was discarded. Proteins were digested with trypsin (Worthington) in 50mM TEAB pH 8.5, at a ratio of 10:1 protein to trypsin, overnight at 37°C. Peptides were eluted with three washes of; first 50 µl 50 mM TEAB, second 50 µl 0.1% formic acid and third 50 µl 50% acetonitrile with 0.1% formic acid. The solution was frozen then dried in a centrifugal concentrator and reconstituted in 15 µl of 0.1% formic acid 2% Acetonitrile.

1 µl of each peptide sample was loaded per LCMS run. Peptides were separated using an UltiMate 3000 RSLCnano HPLC. Samples were first loaded/desalted onto Acclaim PepMap100 C18 LC Column (5 mm Å∼ 0.3 mm i.d., 5 μm, 100 Å, Thermo Fisher Scientific) at a flow rate of 10 μl min−1 maintained at 45°C and then separated on a 75 μm x 75cm C18 column (Thermo EasySpray - C18 2 µm) with integrated emitter using a 60 minutes nonlinear gradient from 92.5% A (0.1% FA in 3% DMSO) and 7.5% B (0.1% FA in 80% ACN 3% DMSO), to 40% B, at a flow rate of 150 nl min−1. The eluent was directed to an Thermo Q Exactive HF mass spectrometer through the EasySpray source at a temperature of 300°C, spray voltage 1500 V. The total LCMS run time was 120 minutes. Orbitrap full scan resolution was 120,000, ACG Target 5e6, maximum injection time 100 ms, scan range 375-1300 m/z. DIA MS/MS were acquired with 15 m/z windows covering 328.5-1251m/z, at 30000 resolution, maximum injection time of 100ms, with ACG target set to 3e6, and normalized collision energy level of 27.

The acquired data has been analysed in DIA-NN (version 1.8)(*62*) against *Clostridium perfringens* proteome sequence database (Uniprot UP000000818, version from 2024/07/16) combined with common Repository of Adventitious Proteins (cRAP), Fragment m/z: 300-1800, enzyme: Trypsin, allowed missed-cleavages: 2, peptide length: 6-30, precursor m/z 300-1250, precursor charge: 2-4, Fixed modifications: carbamidomethylation(C), Variable modifications:Oxidation(M), Acetylation(N-term). The normalised data was then analysed using “Limma” (version 3.60.4)(*63*) in R (version 4.4.0)(*58*). Supernatant and pellet samples were analysed separately. Proteins were deemed significant when associated to an absolute log2 fold change value >1 and adjusted P value <0.05.

### HMO and lactose quantification in bacterial culture supernatants

HMOs and lactose were measured following the method previously published by Molnar-Gabor et al. (2024)(*64*). In brief, this method uses labelling by reductive amination, with 4-aminobenzoic acid ethyl ester (benzocaine) as the labelling reagent and picoline borane as the reducing agent, then applies high-performance liquid chromatography (HPLC) separation with UV detection.

### Fucose, LNB, sialic acid quantification in bacterial culture supernatants

Fucose, LNB, and free sialic acid were measured using HPLC-MS. Samples were diluted tenfold in 1:1 acetonitrile-water, and 50 µl internal standard solution (4 g/l sucrose) was added. A seven-point calibration curve of all three compounds was used. Ultra-high performance liquid chromatography (UHPLC; Thermo Fisher Scientific Ultimate 3000) was used with a binary pump coupled to a Bruker microTOF Q-TOF mass spectrometer. A Thermo Fisher Scientific Accucore 150-Amide-HILIC analytical column was used (dimensions 150 × 3.0 mm, 2.6 μm particle size). A binary gradient of acetonitrile (eluent A) and 50 mM ammonium formate adjusted to pH 3 (eluent B) was used at a flow rate of 0.8 ml/min. The initial composition was 75% A and 25% B, changed to 70% A and 30% B over 5 minutes, then to 60% A and 40% B in 0.2 minutes. This was held for 2.8 minutes to elute any potentially present larger oligosaccharides, then changed to 75% A and 25% B in 0.1 minutes and equilibrated for 3 minutes before the next injection. The MS was used in negative mode, and extracted ion chromatograms at m/z 341 (internal standard, [M-H]^−^), m/z 209 (fucose, [M+HCOOH-H]^−^), m/z 428 (LNB, [M+HCOOH-H]^−^), and m/z 308 (sialic acid, [M-H]^−^) were used for integration, selected based on the mass spectra recorded in the calibration solutions. A quadratic calibration curve with internal standard was used.

### SCFA profiling of *Clostridium* and *Bifidobacterium* culture supernatants and data analysis

For isolates found to grow on HMOs following the above growth curve protocol, supernatants from wells where bacterial growth was observed were harvested, centrifuged at 5000 G for 5 minutes at 4°C to remove any bacteria and the supernatants collected. Quantitative measurement of SCFAs (acetic acid, propionic acid, butyric acid, isobutyric acid, valeric acid, isovaleric acid and hexanoic acid) in the collected samples was then performed by Creative Proteomics, USA, using a gas chromatography-mass spectrometry (GC-MS) method. Samples were diluted in water containing labelled internal standards for each chain length (C2-C6). The free short chain fatty acids were derivatized using methyl chloroformate in 1-propanol yielding propyl esters before subsequent liquid-liquid extraction into hexane and analysis on a SLB-5ms (30 × 0.25 mm × 1.0 µm) column and detection using GC-EI-MS in SIMmode. The analytes were quantified using 8-point calibration curves. For analysis of the data, the raw SCFA concentrations for each strain were divided by the maximum OD600 measured during culture of the strain as part of the growth curve protocol described above, giving growth adjusted concentrations. These values were then averaged for each species. Boxplots of these data were then created in R (version 4.0.4)(*58*), using the ggplot2 package. Between species comparisons for mean SCFA growth adjusted concentrations were performed using an ANOVA followed by Tukey’s honestly significant different (HSD) test in R(*58*), with p <0.05 set at the threshold for statistical significance.

### Preparation of bacterial cell free supernatants

A loopful of a glycerol stock was streaked out across an agar plate of the isolate’s preferred medium and incubated anaerobically at 37°C overnight. A single colony was then picked from this plate, transferred to 5 ml broth based on the isolate’s preferred medium and incubated overnight. Two 5 ml aliquots of ZMB1 were prepared. Glucose was added to 10 mg/mL to one aliquot, while the HMOs known to sustain growth of the isolate were added to the second aliquot, to a total concentration of 10 mg/ml. Due to insufficient material for DSLNT this HMO could not be included in the medium. The OD600 of the overnight growth of the isolate was measured and the volume required to dilute to OD 0.05 in 5 ml of ZMB1 was calculated. Two aliquots of the calculated volume of the culture were then centrifuged at 5000 G for 5 minutes at 4°C, and the pellet was resuspended in 500 μl of anaerobic PBS. Centrifugation was repeated and the pellets then resuspended in 250 μl of ZMB1 + glucose and ZMB1 + HMOs respectively. The culture aliquots were then added to the remaining 4.75 ml of ZMB1 + glucose and ZMB1 + HMOs. The two cultures were then incubated anaerobically, at 37°C, with shaking at 130 rpm, for 2 days. At the end of the incubation period, the two cultures were centrifuged at 5000 G for 10 minutes at 4°C, and the supernatants transferred to new tubes in the anaerobic chamber. These supernatants were then filter sterilised with Merck Millex™-GP Sterile 0.22 μm syringe filters, producing cell free supernatants (CFSs).

### Untargeted metabolomics of cell free supernatants

Metabolomics was performed by Metabolon Inc, as described below.

#### Sample Preparation for untargeted metabolomics

Samples were prepared using the automated MicroLab STAR® system from Hamilton Company. Several recovery standards were added prior to the first step in the extraction process for QC purposes. To remove protein, dissociate small molecules bound to protein or trapped in the precipitated protein matrix, and to recover chemically diverse metabolites, proteins were precipitated with methanol under vigorous shaking for 2 min (Glen Mills GenoGrinder 2000) followed by centrifugation. The resulting extract was divided into multiple fractions: two for analysis by two separate reverse phase (RP)/UPLC-MS/MS methods with positive ion mode electrospray ionization (ESI), one for analysis by RP/UPLC-MS/MS with negative ion mode ESI, one for analysis by HILIC/UPLC-MS/MS with negative ion mode ESI, while the remaining fractions were reserved for backup. Samples were placed briefly on a TurboVap® (Zymark) to remove the organic solvent. The sample extracts were stored overnight under nitrogen before preparation for analysis.

#### QA/QC

Several types of controls were analysed in concert with the experimental samples: a pooled matrix sample generated by taking a small volume of each experimental sample (or alternatively, use of a pool of well-characterized human plasma) served as a technical replicate throughout the data set; extracted water samples served as process blanks; and a cocktail of QC standards that were carefully chosen not to interfere with the measurement of endogenous compounds were spiked into every analysed sample, allowed instrument performance monitoring and aided chromatographic alignment. Instrument variability was determined by calculating the median relative standard deviation (RSD) for the standards that were added to each sample prior to injection into the mass spectrometers. Overall process variability was determined by calculating the median RSD for all endogenous metabolites (i.e., non-instrument standards) present in 100% of the pooled matrix samples. Experimental samples were randomized across the platform run with QC samples spaced evenly among the injections.

#### Data generation

All methods utilized a Waters ACQUITY ultra-performance liquid chromatography (UPLC) and a Thermo Scientific Q-Exactive high resolution/accurate mass spectrometer interfaced with a heated electrospray ionization (HESI-II) source and Orbitrap mass analyser operated at 35,000 mass resolution (PMID: 32445384). The dried sample extract were then reconstituted in solvents compatible to each of the four methods. Each reconstitution solvent contained a series of standards at fixed concentrations to ensure injection and chromatographic consistency. One aliquot was analysed using acidic positive ion conditions, chromatographically optimized for more hydrophilic compounds (PosEarly). In this method, the extract was gradient eluted from a C18 column (Waters UPLC BEH C18-2.1×100 mm, 1.7 µm) using water and methanol, containing 0.05% perfluoropentanoic acid (PFPA) and 0.1% formic acid (FA). Another aliquot was also analysed using acidic positive ion conditions, however it was chromatographically optimized for more hydrophobic compounds (PosLate). In this method, the extract was gradient eluted from the same aforementioned C18 column using methanol, acetonitrile, water, 0.05% PFPA and 0.01% FA and was operated at an overall higher organic content. Another aliquot was analysed using basic negative ion optimized conditions using a separate dedicated C18 column (Neg). The basic extracts were gradient eluted from the column using methanol and water, however with 6.5mM Ammonium Bicarbonate at pH 8. The fourth aliquot was analysed via negative ionization following elution from a HILIC column (Waters UPLC BEH Amide 2.1×150 mm, 1.7 µm) using a gradient consisting of water and acetonitrile with 10mM Ammonium Formate, pH 10.8 (HILIC). The MS analysis alternated between MS and data-dependent MSn scans using dynamic exclusion. The scan range varied slightly between methods but covered 70-1000 m/z. Raw data files are archived and extracted as described below.

#### Bioinformatics

Raw data was extracted, peak-identified and QC processed using a combination of Metabolon developed software services (applications). Each of these services perform a specific task independently, and they communicate/coordinate with each other using industry-standard protocols. Compounds were identified by comparison to library entries of purified standards or recurrent unknown entities. Metabolon maintains a library based on authenticated standards that contains the retention time/index (RI), mass to charge ratio (m/z), and fragmentation data on all molecules present in the library. Furthermore, biochemical identifications are based on three criteria: retention index within a narrow RI window of the proposed identification, accurate mass match to the library +/− 10 ppm, and the MS/MS forward and reverse scores between the experimental data and authentic standards. The MS/MS scores are based on a comparison of the ions present in the experimental spectrum to the ions present in the library spectrum. While there may be similarities between molecules based on one of these factors, the use of all three data points is utilized to distinguish and differentiate biochemicals. More than 5,400 commercially available purified or in-house synthesized standard compounds have been acquired and analysed on all platforms for determination of their analytical characteristics. An additional 7000 mass spectral entries have been created for structurally unnamed biochemicals, which have been identified by virtue of their recurrent nature (both chromatographic and mass spectral). These compounds have the potential to be identified by future acquisition of a matching purified standard or by classical structural analysis. Metabolon continuously adds biologically-relevant compounds to its chemical library to further enhance its level of Tier 1 metabolite identifications.

#### Compound Quality Control

A variety of curation procedures were carried out to ensure that a high-quality data set was made available for statistical analysis and data interpretation. The QC and curation processes were designed to ensure accurate and consistent identification of true chemical entities, and to remove or correct those representing system artifacts, mis-assignments, mis-integration and background noise.

Metabolon data analysts use proprietary visualization and interpretation software to confirm the consistency of peak identification and integration among the various samples.

#### Metabolite Quantification and Data Normalisation

Peaks were quantified using area-under-the-curve. For studies spanning multiple days, a data normalization step was performed to correct variation resulting from instrument inter-day tuning differences. Essentially, each compound was corrected in run-day blocks by registering the medians to equal one (1.00) and normalizing each data point proportionately (termed the “block correction”). For studies that did not require more than one day of analysis, no normalization is necessary, other than for purposes of data visualization. In certain instances, biochemical data may have been normalized to an additional factor (e.g., cell counts, total protein as determined by Bradford assay, osmolality, etc.) to account for differences in metabolite levels due to differences in the amount of material present in each sample.

### Metabolomics data analysis

Following receipt of data from Metabolon Inc., imputation was performed for missing values in the median normalised data. Imputed values were calculated per metabolite by identifying the lowest value measured for each metabolite and dividing it by four. All data were then transformed using the natural logarithm. All subsequent analyses were then performed on these transformed data. Per strain, the metabolomes of glucose and HMO-derived CFSs were compared by PERMANOVA, using the R package Vegan (version 2.6.8)(*65*), with method set to Euclidean distance. Differential abundance analysis comparing metabolite levels in CFSs to those in ZMB1 medium were performed using Limma (version 3.56.2)(*63*), with thresholds set at log_2_(fold change) +/−1 and adjusted *P* value <0.05. *P* values were adjusted using the Benjamini-Hochberg method. Statistical comparisons of metabolite levels between species were performed using ANOVA, followed by Tukey’s HSD test. P <0.05 was set as the threshold for statistical significance.

### Cell free supernatant activity assay against pathobionts and bifidobacteria

A glycerol stock of the isolate being tested was streaked out on an agar plate of the isolate’s preferred medium and incubated anaerobically at 37°C overnight. Single colonies were then picked from the plate, transferred to 5 ml broth based on the isolate’s preferred medium and incubated anaerobically at 37°C overnight. On the same day, a fresh batch of ZMB1 medium was prepared and split into three aliquots. One aliquot was supplemented with glucose to a concentration of 10 mg/ml, the second with glucose to a concentration of 20 mg/ml and the third left as the base medium. All media aliquots were then left in the anaerobic chamber overnight to remove any oxygen. The next day, the OD600 of the bacterial culture was measured and volume required to dilute it to an OD600 of 0.2 in 3 ml ZMB1 + glucose (10 mg/ml) was calculated. The required volume of the culture was then centrifuged at 5000 G for 5 minutes at 4°C and the supernatant removed. The pellet was then resuspended in 1 ml anaerobic PBS and centrifuged again as before, with the supernatants removed. The pellet was then resuspended in 3 ml ZMB1 + glucose (10 mg/ml). The culture was then incubated anaerobically at 37°C, with shaking at 110 rpm, until it reached mid exponential growth phase. At this point, the OD600 was measured and volume required to dilute to an OD600 of 0.1 in 10 ml ZMB1 + glucose (20 mg/ml) was calculated. The required volume of the cultures was then centrifuged at 5000 G for 5 minutes at 4°C and the supernatant removed. The pellet was resuspended in 10 ml ZMB1 + glucose (20 mg/ml). 100 μl of this isolate culture was then added to the wells of a 96-well plate. Glucose-based CFSs (see ‘Preparation of cell free supernatants’) were used for these assays and were thawed under anaerobic conditions. 100 μl of each CFS being tested was added to individual wells. If required for the experiment, CFS pH was adjusted from acidic to neutral using concentrated NaOH, with pH measured using a Thermo Scientific Orion ROSS Ultra pH electrode. Additionally, a SCFA mixture containing acetate and butyrate diluted in ZMB1 was prepared, with concentrations of each based on the average concentrations detected in all *Clostridium* samples during SCFA profiling by Creative Proteomics (acetate: 122.3 mM, butyrate: 47.6 mM). This SCFA mix was split into three aliquots, and each adjusted to pH 4, 6 and 7, using either concentrated HCl or concentrated NaOH as required. 100 μl of SCFA mix at each pH was added to individual wells on the assay plate. Three aliquots of ZMB1 base medium were also adjusted to pH 4, 6 and 7 and 100 μl of each added to individual wells. All conditions were set up in triplicate on the plate. Following this set up, the minimum concentration of glucose in each well was 10 mg/ml. The plate was then shaken at 100 rpm for 1 minute. The lid was then removed, and the plate sealed with a Diversified Biotech Breathe-Easy plate seal. The sealed plate was then placed in a Cerillo Stratus plate reader and the assay run for up to three days, with readings taken at 600 nm, every three minutes. For each test, area under the curve (AUC) for time vs. OD600 was calculated, using a trapezoidal model with the trapz() function from the caTools package (version 1.18.2)(*66*). Statistical comparisons were performed using an ANOVA, followed by Dunnett’s Test comparing each test to the AUC for the growth of each strain with pH7 ZMB1 medium added. The threshold for significance was set at *P* <0.05. For plotting using the package pheatmap (version 1.0.12)(*67*), the AUC for the growth of each strain with pH7 ZMB1 medium added was set as a 100% growth reference for all other conditions and the percentages of this control AUC calculated accordingly. Selected growth curves were generated by plotting mean OD600, calculated from three replicates.

### MTS assay to measure viability of Caco-2 cells incubated with cell free supernatants

Caco-2 cells were cultured in Advanced DMEM, supplemented with FBS to 10% and GlutaMAX. For the assay, the cells were seeded at a density of 5000 cells per well, in a volume of 200 μl per well, across 96 well plates and incubated at 37°C and 5% CO_2_ for 24 hours. The old medium was then removed, and the following conditions set up in 100 μl 1) DMEM only control, 2) base ZMB1 medium, at 10%, 25%, 50% and 75% dilutions in DMEM, 3) 11x HMO CFSs in separate wells at 10%, 25%, 50% and 75% dilutions in DMEM, 4) 12x glucose CFSs in separate wells at 10%, 25%, 50% and 75% dilutions in DMEM. All conditions were set up in triplicate. The assays were then incubated at 37°C and 5% CO2 for 24 hours. Medium was then removed from each well and replaced with 100 μl fresh DMEM. MTS reagent is reduced by metabolically active cells into a formazan salt that is soluble in tissue culture medium, giving a detectable colour change which can be quantified by measuring absorbance at 490 nm using a plate reader. The measured absorbance at 490 nm corresponds to the quantity of formazan produced, which in turn corresponds to the number of living cells in the culture. Thus, the measured absorbance is directly proportional to the number of viable cells and allows the viability of a culture to be compared with others. Fresh MTS reagent was prepared by adding phenazine methosulfate to a concentration of 5% to MTS tetrazolium salt solution. 20 μl of the reagent was then added to each well of the assay plate, which was then incubated at 37°C and 5% CO2 for 4 hours. MTS reagent was also added to triplicate wells containing only DMEM, to act as a blank. The absorbance at 490 nm of each well was then measured. Following blank correction, absorbances were converted to % viabilities, with absorbances for DMEM only controls being used as the denominators in % calculations. DMEM only controls were therefore set as 100% viability and % viabilities for CFS and ZMB1 conditions were calculated relative to those controls. Calculated viabilities for each condition were then averaged. Statistical comparisons were performed at the genus level, with an ANOVA and Tukey HSD test used to compare average % viabilities for *Clostridium* and *Bifidobacterium* CFSs, and blank ZMB1 medium, across each concentration tested. P <0.05 was set as the threshold for statistical significance.

### Organoid media production

Organoid media were made as previously described by Stewart et al. (2020)(*68*).

Complete Media Growth Factor negative (CMGF−): 500 ml Advanced DMEM/F12, 5 ml GlutaMAX 100×, 5 ml HEPES 1 M.

Complete Media Growth Factor positive (CMGF+) in a volume of 500 ml: 78 ml CMGF−, 250 ml Wnt3A-conditioned media produced from ATCC CRL-2647 cells (ATCC), 100 ml R-spondin-conditioned media produced from R-Spondin1 expressing 293T cells (Merck), 50 ml Noggin-conditioned media produced from 293-Noggin cells(*69*), 10 ml B27 (50×), 5 ml N2 (100×), 5 ml nicotinamide (10 mM), 1 ml N Acetylcysteine (1 mM), 500 μl Gastrin (10 nM), 500 μl A83 (500 nM), 166 μl SB202190 (10 μM), 50 μl EGF (50 ng/ml). High Wnt medium in a volume of 500 ml: 250 ml CMGF+, 250 ml Wnt3A-conditioned media.

Differentiation (DIF) medium in a volume of 500 ml: 458 ml CMGF−, 25 ml Noggin conditioned media, 10 ml B27 (50×), 5 ml N2 (100×), 1 ml N Acetylcysteine (500 mM), 500 μl Gastrin (100 μM), 500 μl A83 (500 μM), 50 μl EGF (500 μg/ml).

### Establishment and culture of preterm infant intestinal-derived organoids

Intestinal epithelial organoid lines were established from intestinal crypts isolated from preterm neonate tissue, as described by Stewart et al. (2020)(*68*). Briefly, tissue was minced, washed with chelating solution, antifungals and antibiotics, and crypt cells extracted by gentle shaking in EDTA. Extracted cells were then suspended within phenol-red free and growth factor reduced Matrigel basement membrane matrix (Corning), which was then pipetted as small dots into the wells of a 24 well plate, at a maximum of three dots per well. The Matrigel was set by incubating at 37°C for 30 minutes. Following polymerization of the Matrigel, 500 μl High Wnt growth medium was added to each well and the organoids left to grow from the crypt cells, with incubation at 37°C and 5% CO_2_. Media was changed for fresh every Monday, Wednesday and Friday.

Organoids were passaged by removing spent medium and adding 300 μl 0.05% Trypsin-EDTA (Gibco) per well. The typsin was then pipetted up and down to break up the Matrigel and suspend the organoids.

Organoids were incubated in trypsin at 37°C and 5% CO_2_ for 5 minutes, with 350 μl fetal bovine serum (Merck) then added to stop trypsinisation. Following centrifugation at 363 G for 5 minutes, supernatant was removed, and the organoids resuspended in ice cold Matrigel, the volume of which depended on the number of wells being seeded. The Matrigel was then pipetted as small dots into the wells of a 24 well plate, at a maximum of three dots per well and set by incubating at 37°C for 30 minutes. Following polymerization of the Matrigel, 500 μl High Wnt growth medium was added to each well and the organoids left to grow, with incubation at 37°C and 5% CO_2_. All organoid experiments were conducted with cell line NCL 27, within passages 10-15. This line was established from a patient born at 24 weeks gestation, using ileum tissue salvaged from surgery performed on day of life 10 due to the development of necrotising enterocolitis.

### Organoid monolayer co-culture with bacterial CFSs and inflammatory stimuli

Three dimensional organoids were processed into two dimensional monolayers in 6.5 mm Transwells (Corning), following the protocol described by Fofanova et al. (2024)(*35*). Briefly, 3D organoid cultures were processed into single cell suspensions by washing with EDTA, trypsinisation (0.05% trypsin-EDTA) and passage through a 40 μm nylon cell strainer (Corning). Cells were then resuspended in CMGF+ growth medium (1.8×10^6^ cells per ml) and seeded onto Transwells, pre-coated with diluted (1:40 in cold PBS) Matrigel. Growth medium was replaced with DIF medium after 2-3 days, once trans-epithelial electrical resistances (TEERs) were near 300 Ω, indicating monolayer confluence. Monolayers were differentiated for four days prior to experiments. Monolayers were then placed into the anaerobic co-culture system(*35*), which was in turn placed into an anaerobic chamber. Oxygen is flowed through the base of the co-culture unit, maintaining aerobic conditions on the basolateral sides of the monolayers, while the apical sides remain under the anaerobic conditions of the chamber. Thus, the physiological oxygen gradient of the gut epithelium is recreated in this model. The inflammatory stimuli used were lipopolysaccharide (LPS) from *Escherichia coli* 0111:B4 (InvivoGen) and flagellin from *Salmonella typhimurium* (InvivoGen). CFSs were prepared as described above. The following conditions were set up in triplicate across the monolayers: 1) No stimuli, no CFS (‘Control’), 2) + apical CFS to 25% v/v (‘CFS (25%)’), 3) + apical LPS (100 ng/ml) + basolateral flagellin (100 ng/ml) (‘Stimuli’), 4) + apical CFS to 25% v/v + apical LPS (100 ng/ml) + basolateral flagellin (100 ng/ml) (‘CFS (25%) and Stimuli’). The experiment was run for three hours. The TEERs of the monolayers were measured before the start and at the end of the experiment. End point TEERs were subtracted from start point TEERs to give change in TEER during experiment. Outliers were identified by calculating robust/modified z scores for each data point. This method uses the median to calculate median absolute deviation. Any data point with a robust z score greater than +3.5 or less than −3.5 was removed from the dataset. Mean change in TEER for each condition was then calculated. Statistical comparisons of change in TEER between conditions were performed with an ANOVA and Tukey HSD test. Conditions with matching letters were not significantly different. Furthermore, after the incubation, apical and basolateral supernatants were harvested for cytokine assays.

### Cytokine assays

Secreted interleukin-8 (IL-8) was measured using a DuoSet ELISA kit, following the manufacturer’s protocol. All other cytokines were measured using custom U-Plex assays (Meso Scale Discovery Inc.), following the manufacturer’s protocol. Outliers were identified by calculating robust/modified z scores for each data point. This method uses the median to calculate median absolute deviation. Any data point with a robust z score greater than +3.5 or less than −3.5 was removed from the dataset. ANOVA followed by a Dunnett’s test to compare each condition to the untreated control were used to perform statistical analysis of cytokine secretion data. For comparisons of all against all, the ANOVA was followed by Tukey’s HSD test. P <0.05 was set as the threshold for statistical significance.

### Organoid monolayer co-culture with live *C. perfringens* and inflammatory stimuli

Three dimensional organoids were processed into two dimensional monolayers in 6.5 mm Transwells, following the protocol described by Fofanova et al. (2019)(*35*). Monolayers were then placed into the anaerobic co-culture system, which was in turn placed into an anaerobic chamber. Oxygen is flowed through the base of the co-culture unit, maintaining aerobic conditions on the basolateral sides of the monolayers, while the apical sides remain under the anaerobic conditions of the chamber. Thus, the physiological oxygen gradient of the gut epithelium is recreated in this model. The inflammatory stimuli used were LPS from *E. coli* 0111:B4 (InvivoGen) and flagellin from *S. typhimurium* (InvivoGen). ZMB1 medium was prepared a day ahead, mixed 1:1 with organoid DIF medium and this mixture left in the anaerobic chamber overnight. *C. perfringens* AM1 and JC26 were grown overnight in BHI broth. The next day, the OD600 of each culture was measured and volumes required to dilute each to an OD600 of 0.2 in 2 ml ZMB1:DIF were calculated. The required volumes of the cultures were then centrifuged at 5000 G for 5 minutes at 4°C and the supernatants removed. The pellets were then resuspended in 1 ml anaerobic PBS and centrifuged again as before, with the supernatants removed. All pellets were then resuspended in 2 ml ZMB1:DIF. The cultures were then incubated anaerobically at 37°C, with shaking at 110 rpm, until they each reached mid exponential growth phase. The OD600 of each was then measured and volumes required to dilute them to an OD600 of 0.2 in 0.2 ml ZMB1 were calculated. The required volumes of the cultures were then centrifuged at 5000 G for 5 minutes at 4°C and the supernatants removed. The pellets were then resuspended in 0.2 ml ZMB1:DIF. These cultures were then added to the apical side of the monolayers as required. A second culture of JC26 was prepared in the same way but timed to be added to the appropriate monolayers at the end of the first hour of the experiment. The following conditions were set up in triplicate across the monolayers: 1) No stimuli, no bacteria, 2) + AM1, 3) + JC26, 4) + AM1 for one hour, then add JC26, 5) No treatment for one hour then add apical LPS (100 ng/ml) and basolateral flagellin (100 ng/ml), 6) + AM1 for one hour, then add apical LPS (100 ng/ml) and basolateral flagellin (100 ng/ml). The ZMB1:DIF mixture was used as the apical medium across all conditions.

The experiment was run for 3 hours. The trans-epithelial electrical resistances (TEERs) of the monolayers were measured with an epithelial Ohm meter before the start and at the end of the experiment. End point TEERs were subtracted from start point TEERs to give change in TEER during experiment. Mean change in TEER for each condition was then calculated. Outliers were identified by calculating robust/modified z scores for each data point. This method uses the median to calculate median absolute deviation. Any data point with a robust z score greater than +3.5 or less than −3.5 was removed from the dataset. Statistical comparisons of change in TEER between conditions were performed with an ANOVA and Tukey HSD test. P <0.05 was set as the threshold for statistical significance. Conditions with matching letters were not significantly different. Furthermore, after the incubation, apical and basolateral supernatants were harvested for cytokine assays.

### Proteomics on *C. perfringens* AM1 and JC26 cell free supernatants

CFSs were generated as described above. Proteins were purified as described in section ‘Proteomics of *C. perfringens* AM1 grown on specific HMOs’. An equivalent of 5 μg of total protein was then used for digestion. Digested protein was then reduced, alkylated, digested and washed as described above, with the exception that they were finally reconstituted in 20 μl of 0.1% formic acid 2% acetonitrile. ∼0.5 μg of each sample was loaded onto Evotips, according to manufacturer instructions and peptides separated on a 8 cm x 100 µm Evosep Endurance C18 column (Evosep, EV1094) using an Evosep One system (Evosep), with a predefined samples per day protocol 60. The 21 minute gradient ran from 0-35% solvent B (solvent A: 0.1 % formic acid in water, solvent B: 0.1 % formic acid in acetonitrile) at 100 nl/min. Through a 20 µm captive spray emitter (Bruker), at 50 °C, the analytes were directed to a timsToF HT mass spectrometer (Bruker Daltonics). The instrument operated in DIA-PASEF mode, acquiring mass and ion mobility ranges of 300-1400 m/z and 0.6-1.4 1/K0, with a total of 16 variable IM-m/z windows with two quadrupole positions per window, designed using py_diAID from a pooled sample subjected to DDA-PASEF(*70*). TIMS ramp and accumulation times were 100 ms, total cycle time was ∼1.8 seconds. Collision energy was applied in a linear fashion, where ion mobility = 0.6-1.6 1/K0, and collision energy = 20 - 59 eV. The acquired data were analysed as described in ‘Proteomics of *C. perfringens* AM1 grown on specific HMOs’.

### Seahorse Mitochondrial Stress Test

The impacts of CFSs on organoid mitochondrial bioenergetic function were tested using the Seahorse XF Cell Mito Stress Test (Agilent Technologies). This is a live cell assay that detects changes in parameters of mitochondrial function through directly measuring the oxygen consumption rate (OCR) of cells in response to the addition of modulators of respiration. The modulators used were oligomycin (2 μM), Carbonyl cyanide-4 (trifluoromethoxy) phenylhydrazone (FCCP) (4 μM), rotenone (0.5 μM) and antimycin A (0.5 μM). Stocks of the modulators were diluted in organoid CMGF-medium for use in this experiment. The Seahorse XFe96 sensor cartridge (Agilent Technologies) containing these modulators was prepared following the manufacturer’s instructions. Organoid monolayers were generated from NCL 27 at passage 12, as described above, with the exception that they were seeded into a Seahorse XFe96/XF Pro Cell Culture Microplate (Agilent Technologies) at a concentration of 0.3×10^6^ cells per mL, 200 μl per well. Prior to seeding, the Seahorse Microplate was coated with 32 μl per well of Matrigel diluted 40:1 in PBS. The PBS was removed prior to monolayer seeding. Monolayers were differentiated for 4 days prior to start of experiment. On the day of the experiment, the DIF medium was removed from the plate and the cells washed twice in 200 μl pre-warmed CMGF-medium. After washing, 135 μl CMGF-medium was added to each well. Each CFS being tested or ZMB1 (base medium) were then added to individual wells, to final concentrations of 25% v/v and giving a total volume of 180 μl, following the manufacturer’s protocol. A combination of LPS (100 ng/ml) and flagellin (100 ng/ml) were also added to a set of wells to create an “inflammatory stimuli” condition. A set of wells containing only CMGF-were also set up, as negative controls. All conditions were set up in triplicate. The plate was then incubated for one hour, at 37°C in a non-CO_2_ incubator, per the manufacturer’s protocol. The plate was then loaded into a Seahorse XF96 Analyser (Agilent Technologies) and the XF Mito Stress Test protocol run. Following instrument calibration, the assay ran for 73 minutes. Raw data was then processed and exported from the associated software Wave (version 2.6.3, Agilent Technologies). Outliers were identified by calculating robust/modified z scores for each data point. This method uses the median to calculate median absolute deviation. Any data point with a robust z score greater than +3.5 or less than −3.5 was removed from the dataset. Statistical analyses of raw OCR values for each parameter measured were performed using an ANOVA, followed by a Dunnett’s test to compare each condition to the negative control. P <0.05 was set as the threshold for statistical significance.

### *C. perfringens* AM1 and JC26 growth curves in ZMB1:DIF medium

Glycerol stocks of AM1 and JC26 were streaked out on BHI agar plates and incubated in an anaerobic chamber at 37°C overnight. Single colonies were then picked from these plates, transferred to 3 ml BHI broth and incubated in the anaerobic chamber at 37°C overnight. A 50:50 mixture of ZMB1 and DIF media was also prepared and incubated in the anaerobic chamber overnight. The next day, the OD600 of each culture was measured and volumes required to dilute each to an OD600 of 0.2 in 5 ml ZMB1:DIF were calculated. The required volumes of the cultures were then centrifuged at 5000 G for 5 minutes at 4°C and the supernatants removed. The pellets were then resuspended in 1 ml anaerobic PBS and centrifuged again as before, with the supernatants removed. All pellets were then resuspended in 5 ml ZMB1:DIF. Three 5 ml cultures each of AM1 and JC26 were set up in this manner. The cultures were then incubated anaerobically at 37°C, with shaking at 110 rpm. The OD600 of all cultures were measured firstly at 60 minutes and then every 30 minutes, up to 180 minutes. Statistical comparisons between the two strains at each timepoint were made using an unpaired t test, with a threshold of p <0.05 set for significance.

## Acknowledgments

Human milk oligosaccharides were kindly donated by dsm-firmenich through the HMO Donation Program. We thank Dóra Molnár-Gábor (dsm-firmenich, Denmark) for running high-performance liquid chromatography of the cell free supernatants. We thank Simon Kroll, Kathleen Sim and Alexander G Shaw (Imperial College London) providing some of the *Clostridium perfringens* isolates presented in Supplementary Figure 1. We further appreciate the support of Jeremy Palmer with the Meso Scale Discovery assays.

## Author contributions

ACM, JAC and CJS conceived and designed the experiments. ACM, JAC and JAD tested bacterial isolate growth on HMOs. GY analysed *Clostridium* prevalence in the MAGPIE dataset. RK and LJH performed phylogenetic and genomic analyses on *Clostridium perfringens* isolates and provided additional bacterial isolates for testing on HMOs. ACM performed the experiments, sample preparation and data analyses to characterise *Clostridium* HMO utilisation genes and proteins. JPRC and AN supported the RNA-Seq and PP and AP performed the proteomics. ML performed the high-performance liquid chromatography of the cell free supernatants. ACM and JAC generated samples for SCFA and metabolomics analysis and JAC analysed the data. JAC performed pathobiont and *Bifidobacterium* growth challenge assays using CFSs. PP and AP performed proteomics on CFSs and JAC analysed the data. JAC performed the MTS assay. ACM and JAC cultured PIOs. JAC performed organoid co-cultures with CFSs and live bacteria including ELISAs and MSD cytokine assays. JAC performed AM1 and JC26 growth curves. JAC, HW, and CAL designed and performed organoid Seahorse assays and JAC analysed the data. JDP performed MALDI-TOF to identify bacterial isolates at the genus or species level. HPB, YS and TDL performed whole genome sequencing of bacterial isolates. LB performed genome assembly and annotation. JEB and NDE oversaw patient sample collection. CJS conceived and supervised the study. JAC, ACM and CJS prepared figures and wrote the paper. All authors approved the final version of the manuscript.

## Data availability

All cytokine data from ELISA and multiplex MSD assays is provided in Supplementary tables 5-11. The RNA-seq data have been deposited in the Sequencing Read Archive (SRA) under study accession number PRJNA1214204. The proteomics datasets are deposited in MassIVE under submission ID MSV000096907.

