## Supplementary material for "Human milk oligosaccharide metabolism by *Clostridium* species suppresses inflammation and pathogen growth": fig S1

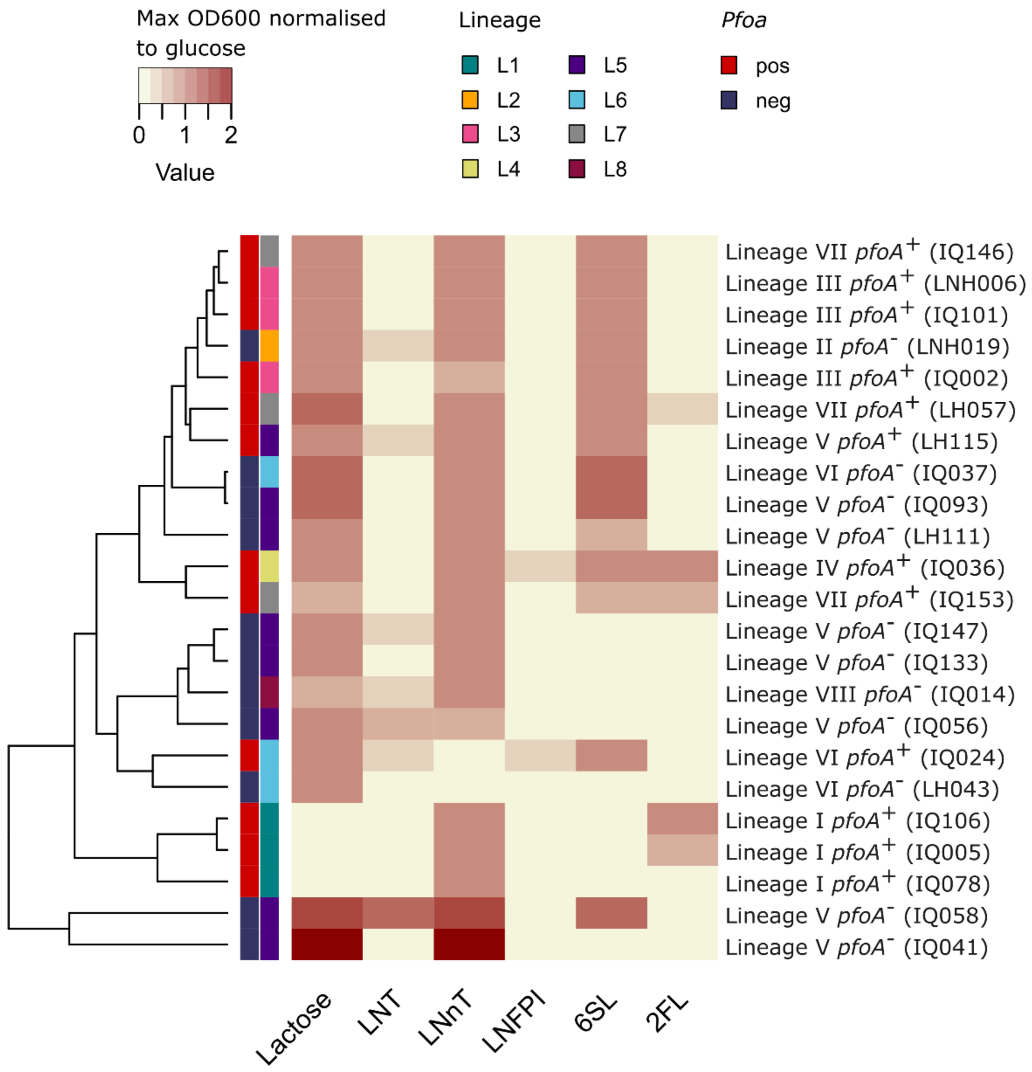


**Figure S1. Growth of *Clostridium perfringens* isolates obtained from Kiu *et al.* (2023)(*21*) on HMOs.** Heatmap representing the growth of 23 bacterial isolates on 6 HMOs and lactose. The values reported represent the maximum OD600 reached normalised to glucose.
LNT, lacto-N-tetraose; LNnT, lacto-N-neotetraose; LNFP I, lacto-N-fucopentaose, 2’FL, 2’ fucosyllacotse.
