## Supplementary material for "Human milk oligosaccharide metabolism by *Clostridium* species suppresses inflammation and pathogen growth": fig S2

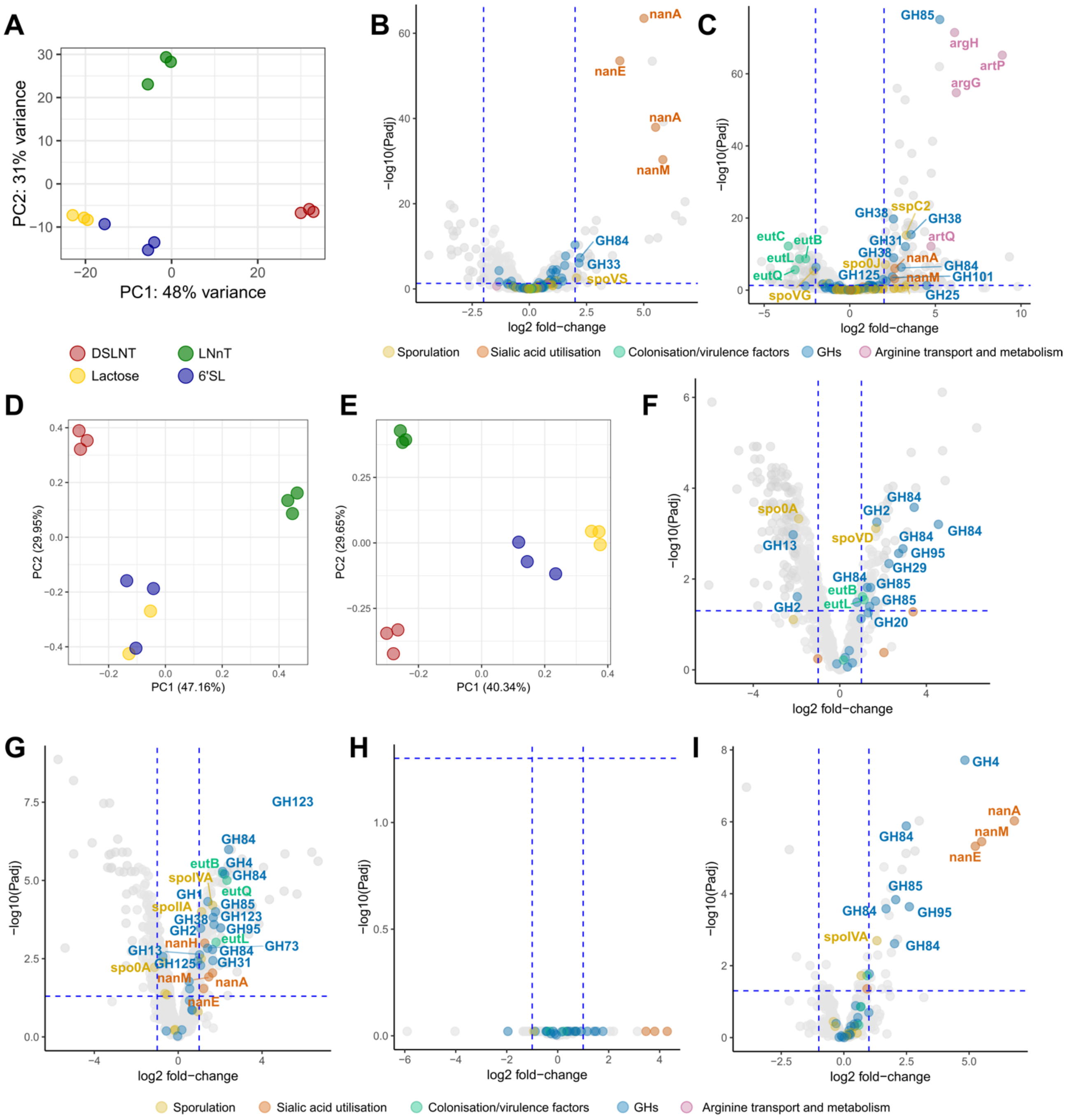


**Figure S2. RNA-seq and proteomics analysis of AM1 grown on DSLNT, LNnT, 6'SL and lactose.** (**A**) Principal Components Analysis (PCA) of RNA-seq data from AM1 grown to exponential phase on DSLNT, LNnT, 6'SL and lactose. (**B-C**) Volcano plot of RNA-seq data for AM1 grown to exponential phase on 6'SL (B) and LNnT (C) compared to lactose. A positive log2 fold-change indicates upregulation on LNnT or 6'SL relative to lactose, while a negative fold change indicates downregulation on LNnt or 6'SL relative to lactose. (**D-E**) PCA of proteomics data from AM1 grown to exponential phase on DSLNT, LNnT, 6'SL and lactose. (**F-G**) Volcano plot of proteomics data from supernatant (F) and pellet (G) of AM1 grown to exponential phase on LNnT compared to lactose. A positive log2 fold-change indicates upregulation on LNnT relative to lactose, while a negative fold change indicates downregulation on LNnt relative to lactose. (**H-I**) Volcano plot of proteomics data from supernatant (F) and pellet (G) of AM1 grown to exponential phase on 6'SL compared to lactose. A positive log2 fold-change indicates upregulation on 6'SL relative to lactose, while a negative fold change indicates downregulation on 6'SL relative to lactose.
LNT, lacto-N-tetraose; LNnT, lacto-n-neotetraose; LNFP I, lacto-N-fucopentaose I; 6'-SL, 6'-sialyllactose; 2'-fucosyllactose,; DSLNT, disialyllacto-N-tetraose; NEC, necrotising enterocolitis; GH, glycoside hydrolase.
