## Supplementary material for "Human milk oligosaccharide metabolism by *Clostridium* species suppresses inflammation and pathogen growth": fig S3

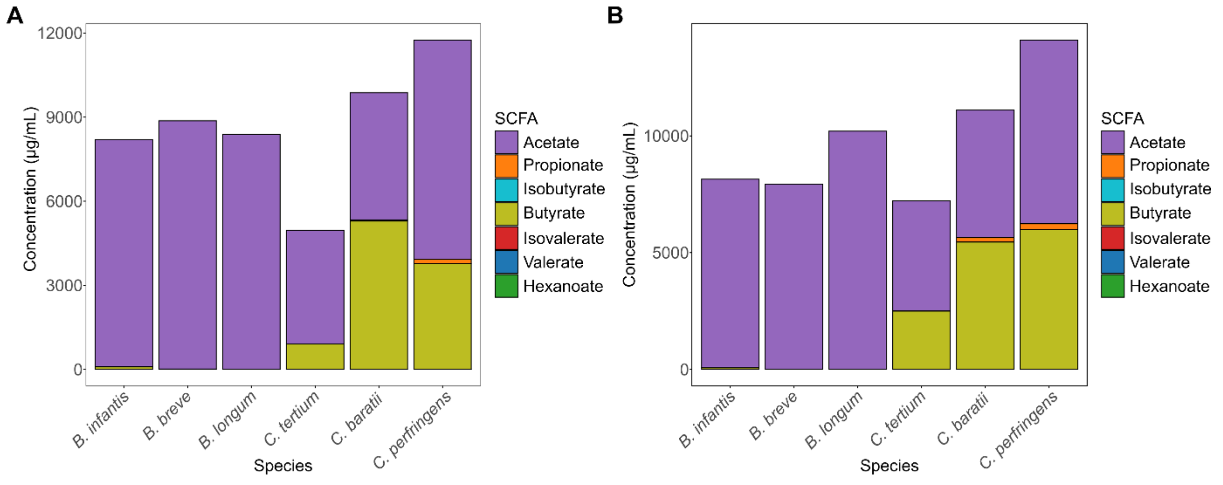


**Figure S3. Total concentration of short chain fatty acids (SCFAs) in culture supernatants of *Bifidobacterium and Clostridium* spp. grown on glucose (A) and lactose (B).**
