## Supplementary material for "Human milk oligosaccharide metabolism by *Clostridium* species suppresses inflammation and pathogen growth": fig S4

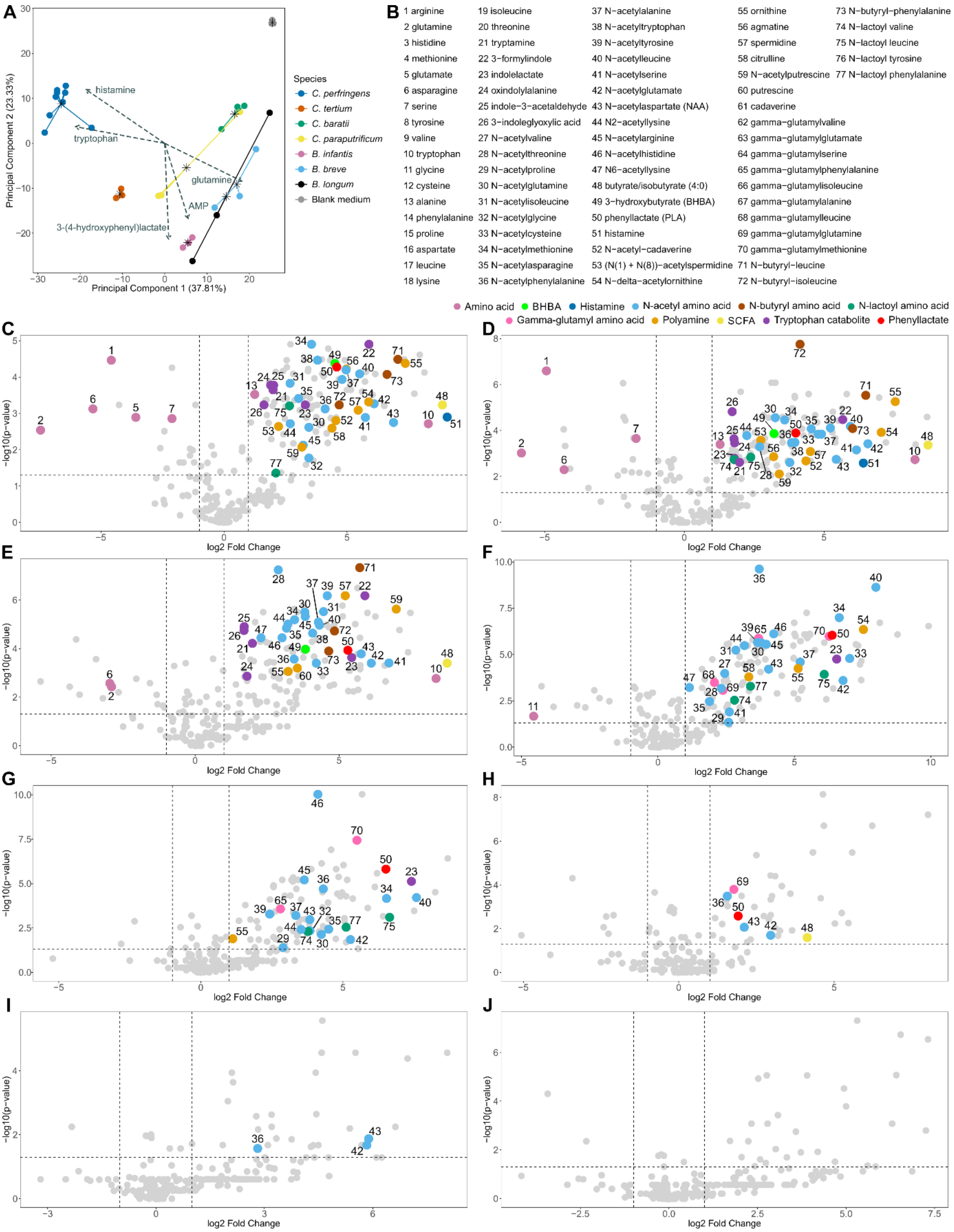


**Figure S4. Untargeted metabolomics analysis of cell free supernatants (CFSs) of isolates grown on a mixture of HMOs.**

(**A**) PCA of untargeted metabolomics data for CFSs generated from bacterial cultures growing on glucose. Arrows indicate the top five metabolites by loadings magnitude. (**B-J**) Volcano plots comparing the metabolites detected in CFSs of isolates growing on a mixture of HMOs compared with blank ZMB1 medium. A positive log2 fold-change indicates production of metabolites by the strain, while a negative fold change indicates metabolite depletion. Metabolites of interest are

highlighted and numbered. (B) Numbered key for labelled metabolites of interest. (C) Volcano plot for CFS of *C. perfringens* pfoA$^-$ AM1 growing on cocktail of the HMOs 6'SL and LNnT, compared with blank ZMB1 medium. (D) Volcano plot for CFS of *C. perfringens* Adult JC36 growing on cocktail of the HMOs 6'SL, LNnT, 2FL, LNT and LNFPI, compared with blank ZMB1 medium. (E) Volcano plot for CFS of *C. tertium* JC25 growing on cocktail of the HMOs LNT and LNnT, compared with blank ZMB1 medium. (F) Volcano plot for CFS of *B. infantis* LB1 growing on cocktail of the HMOs 6'SL, LNnT, 2'FL, LNT and LNFPI, compared with blank ZMB1 medium. (G) Volcano plot for CFS of *B. breve* AM76 growing on cocktail of the HMOs LNT and LNnT, compared with blank ZMB1 medium. (H) Volcano plot for CFS of *C. baratii* AM37 growing on cocktail of the HMOs LNT and LNnT, compared with blank ZMB1 medium. (I) Volcano plot for CFS of *C. paraputrificum* JC53 growing on cocktail of the HMOs LNT and LNnT, compared with blank ZMB1 medium. (J) Volcano plot for CFS of *B. longum* AM7 growing on the HMO LNT, compared with blank ZMB1 medium.
BHBA, beta-hydroxybutyric acid; SCFA, short chain fatty acid.
