## Supplementary material for "Human milk oligosaccharide metabolism by *Clostridium* species suppresses inflammation and pathogen growth": fig S5

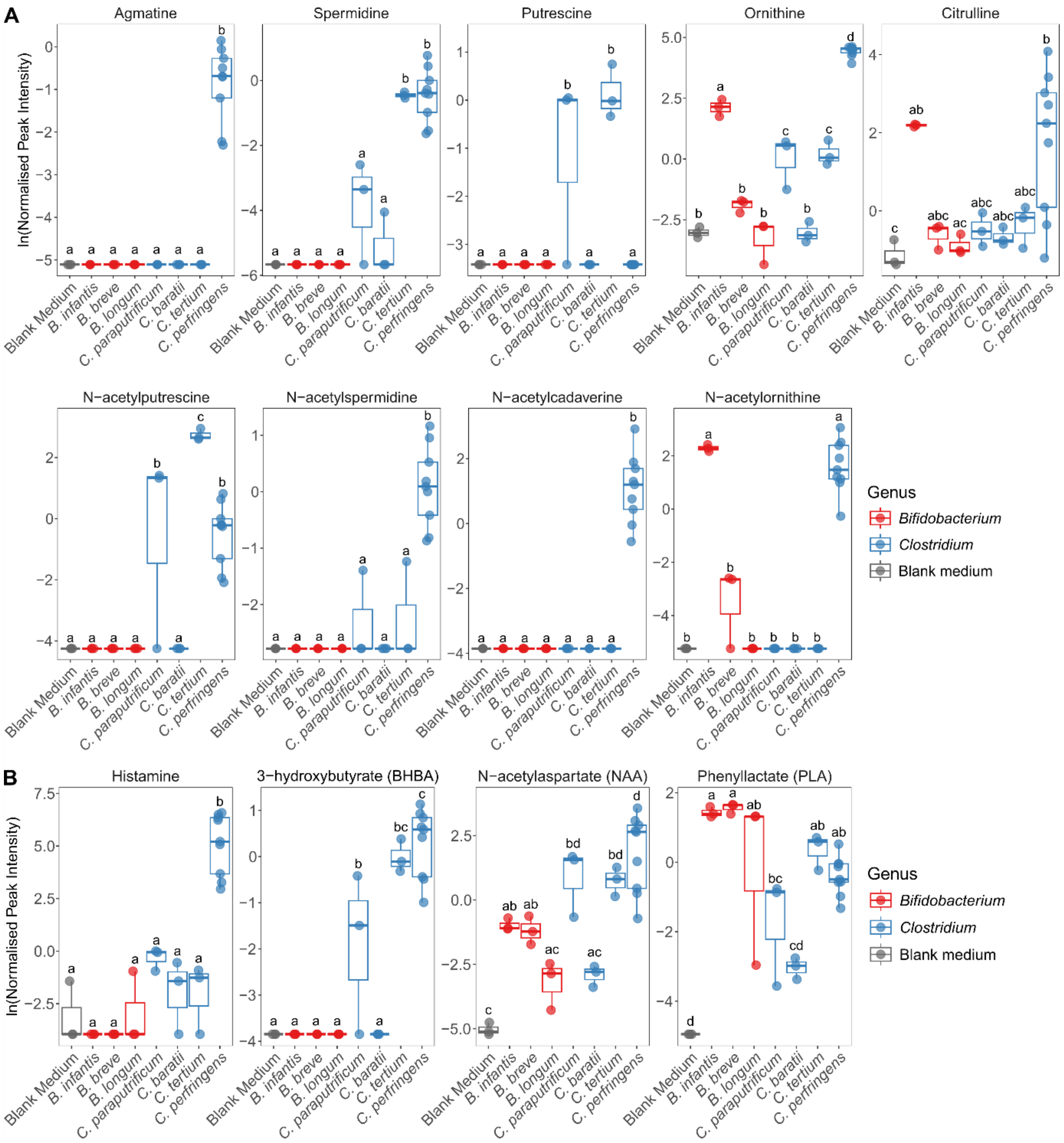


**Figure S5. Levels of metabolites of interest produced by *Bifidobacterium* and *Clostridium* spp. grown on human milk oligosaccharides.**

(**A**) Levels of polyamines produced by *Bifidobacterium* and *Clostridium* spp. during growth on HMOs. Conditions with the same letters are not significantly different. (**B**) Levels of neuromodulatory, immunomodulatory or antimicrobial metabolites produced by *Bifidobacterium* and *Clostridium* spp. during growth on HMOs. Conditions with the same letters are not significantly different.

HMOs, human milk oligosaccharides.
