## Supplementary material for "Human milk oligosaccharide metabolism by *Clostridium* species suppresses inflammation and pathogen growth": fig S6

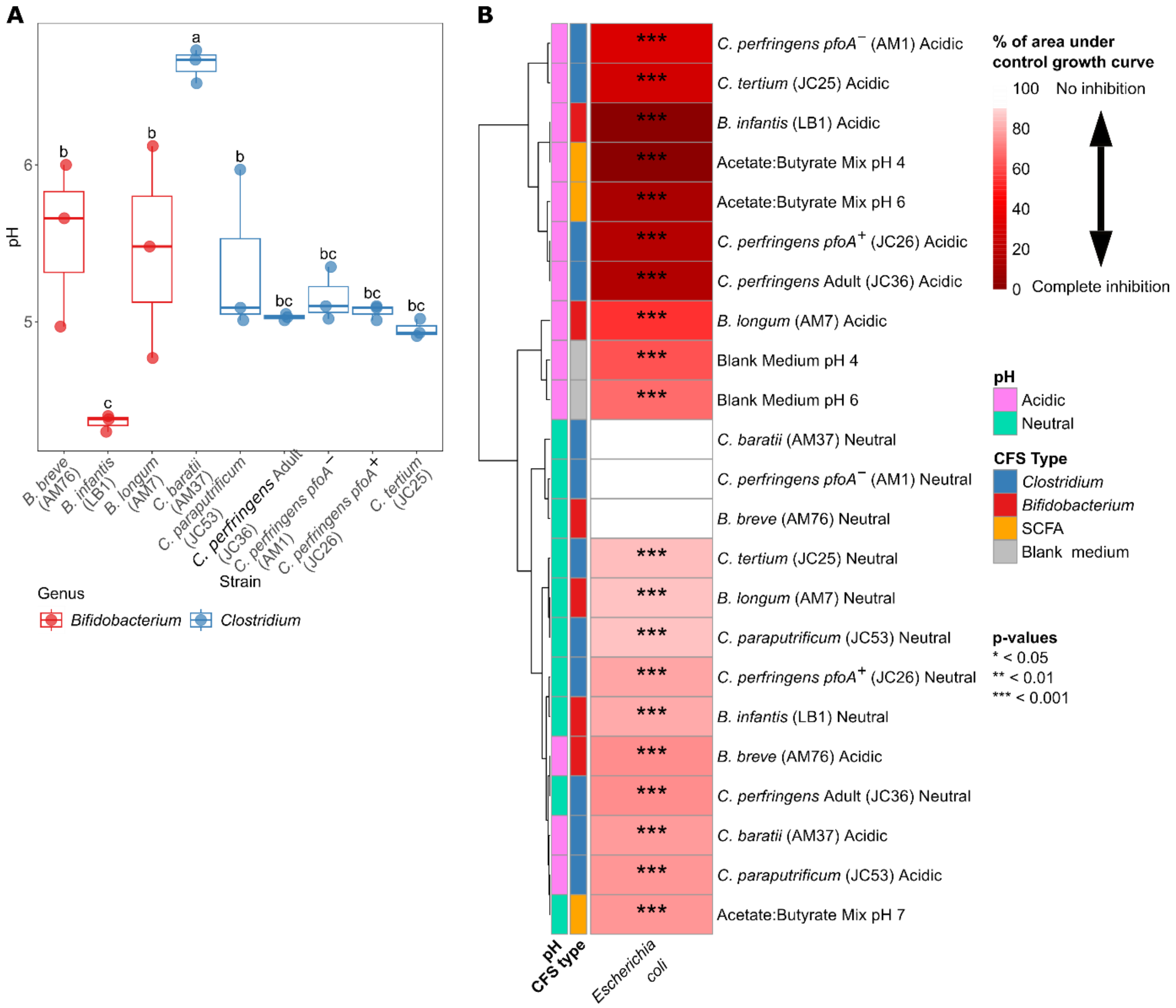


**Figure S6. Effect of pH neutralisation on the antimicrobial activity of all cell free supernatants (CFSs)**

(**A**) Measured pHs of three batches of each CFS generated. Conditions with the same letters are not significantly different. (**B**) Comparison of effect of pH neutralisation on activities of all postbiotics on *E. coli* growth.

SCFA, short chain fatty acid.
