## Supplementary material for "Human milk oligosaccharide metabolism by *Clostridium* species suppresses inflammation and pathogen growth": fig S7

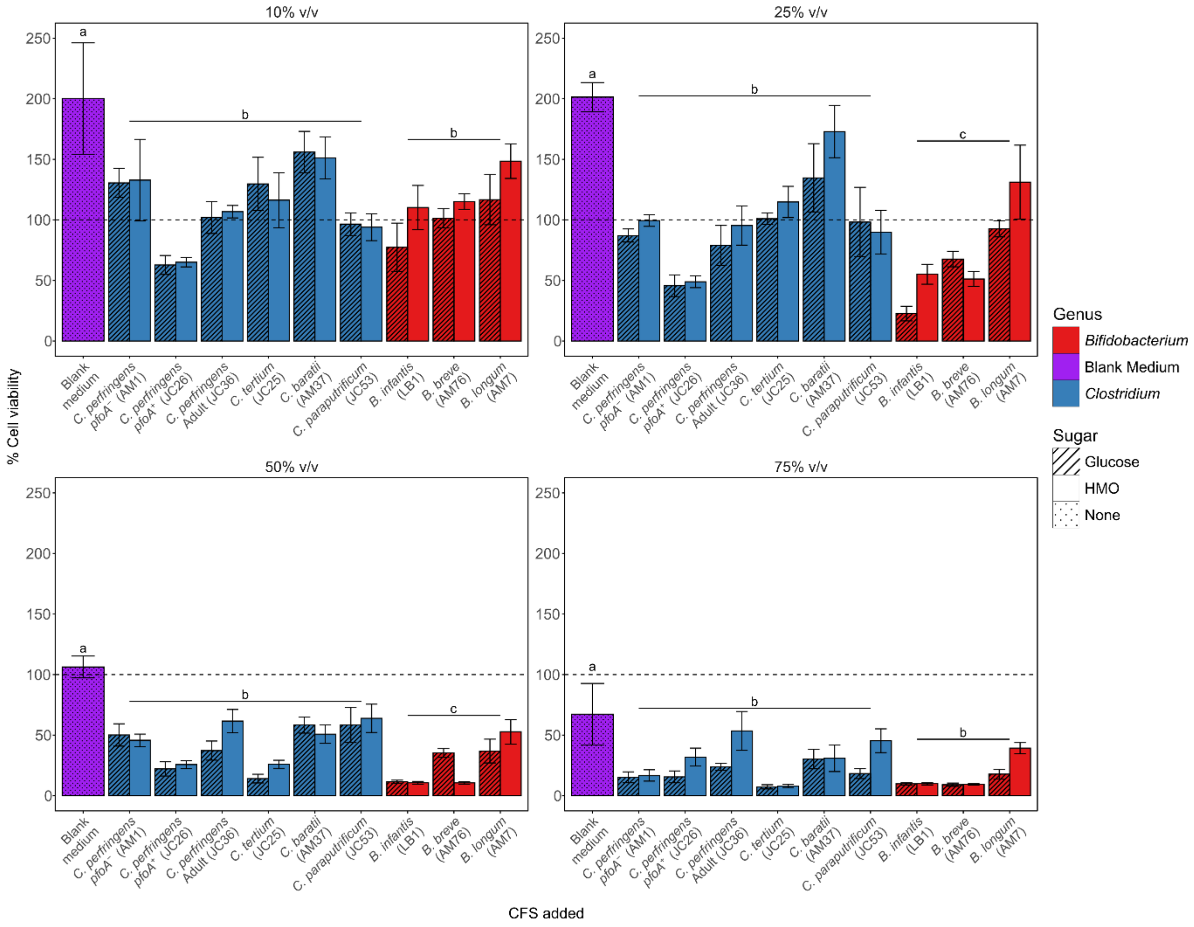


**Figure S7. MTS tetrazolium dye cell viability assay data showing toxicity of cell free supernatants to Caco-2 cells.** Conditions with the same letters are not significantly different. Statistical comparisons were performed using average genus values. Values are shown as % of the viability measured for Caco-2 cells incubated in regular DMEM.

CFS, cell free supernatant; HMO, human milk oligosaccharide.
