## Supplementary material for "Human milk oligosaccharide metabolism by *Clostridium* species suppresses inflammation and pathogen growth": fig S8

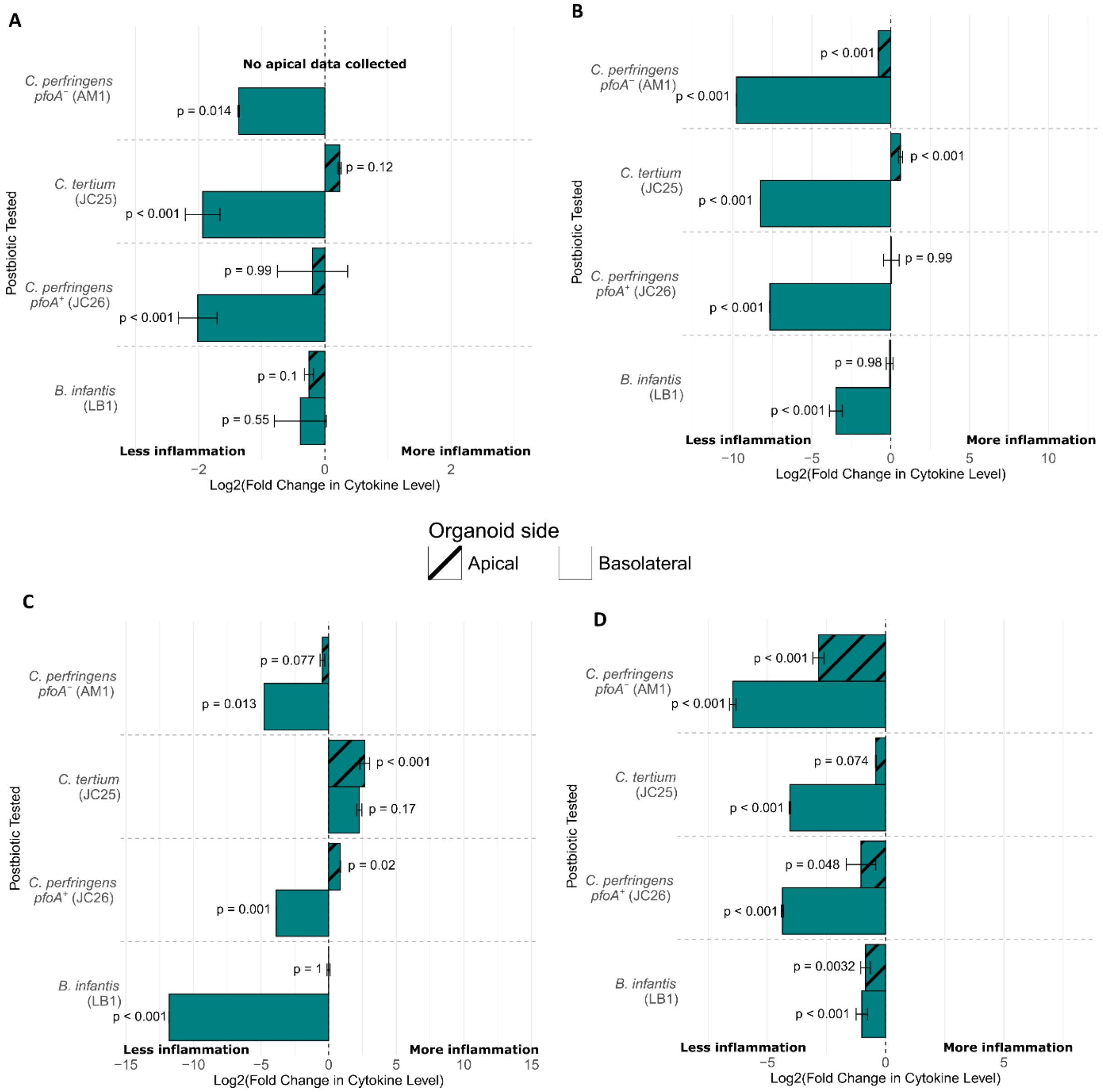


**Figure S8. Changes in organoid monolayer apical and basolateral cytokine secretion during combined treatment with both 25% v/v cell free supernatants and inflammatory stimuli.** Data has been converted to fold change vs "Stimuli" condition and log2 transformed for plotting. P values are for the differences between the unprocessed detected cytokine levels. (**A**) CXCL5 (**B**) CCL7 (**C**) TNF-α (**D**) CCL2
