## Supplementary material for "Human milk oligosaccharide metabolism by *Clostridium* species suppresses inflammation and pathogen growth": fig S9

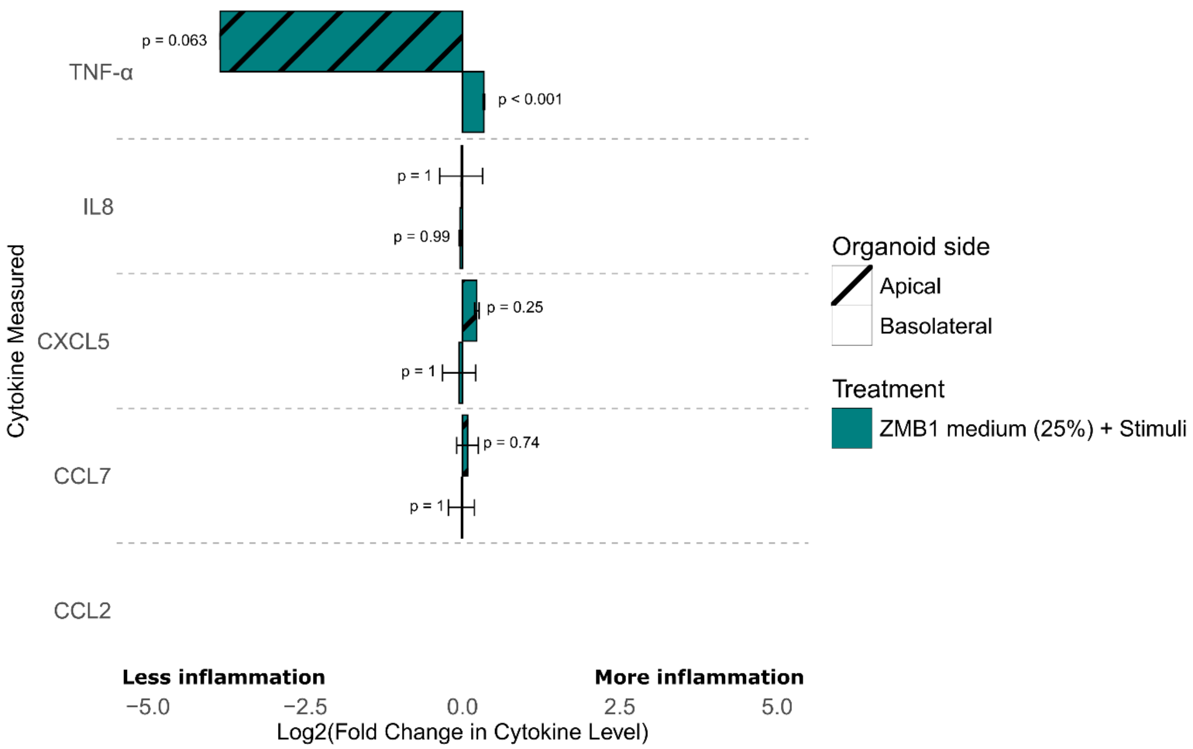


**Figure S9. Changes in organoid monolayer apical and basolateral cytokine secretion during treatment with ZMB1 bacterial growth medium only, at 25% v/v concentration**. Data has been converted to fold change vs "Stimuli" condition and log2 transformed for plotting. P values are for the differences between the unprocessed detected cytokine levels.
