## Supplementary material for "Human milk oligosaccharide metabolism by *Clostridium* species suppresses inflammation and pathogen growth": fig S10

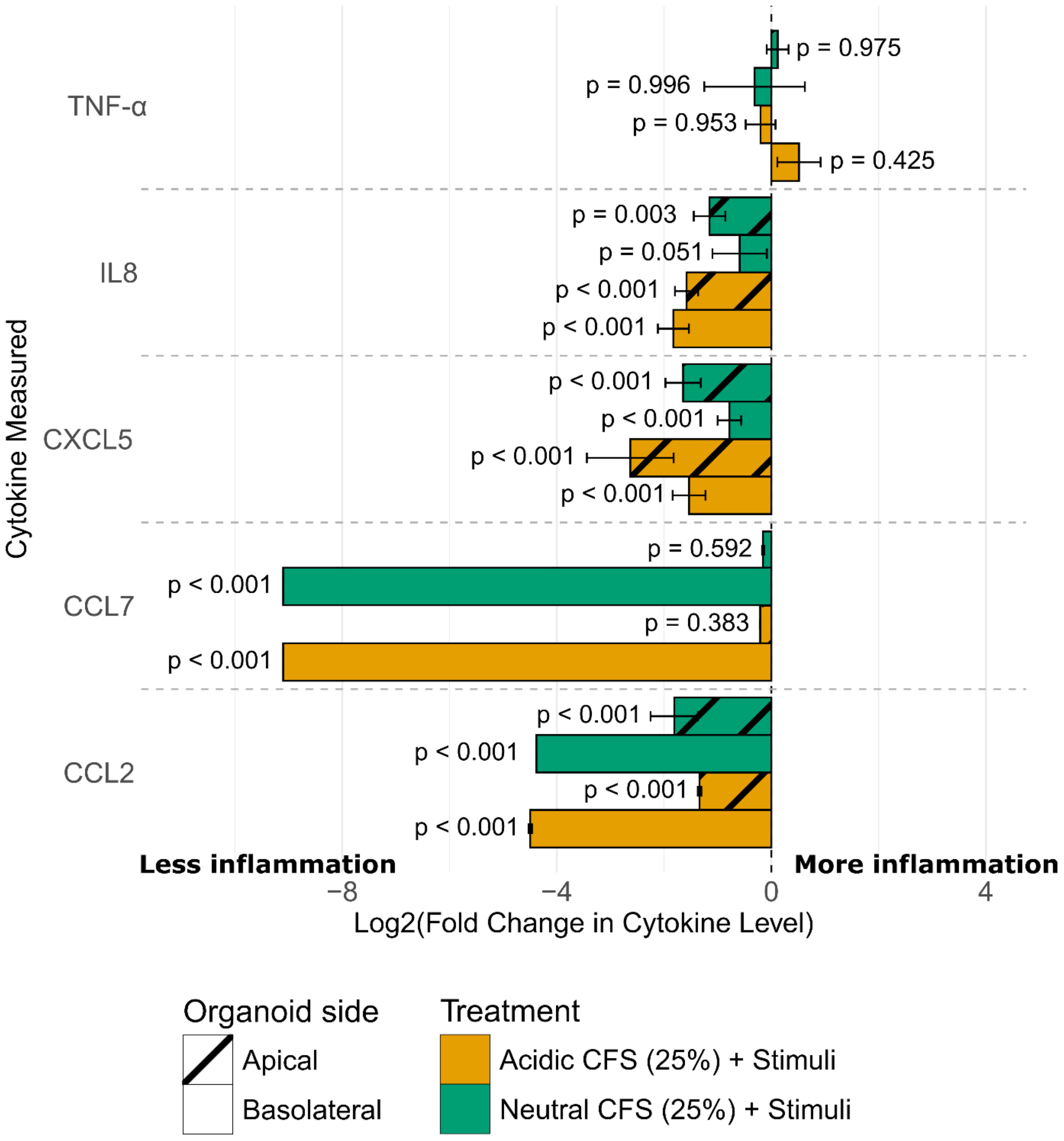


**Figure S10. Changes in organoid monolayer apical and basolateral cytokine secretion during treatment with acidic and neutral pH *C. perfringens* AM1 cell free supernatant (CFS), at 25% v/v concentration.** Data has been converted to fold change vs "Stimuli" condition and log2 transformed for plotting. P values are for the differences between the unprocessed detected cytokine levels.

CFS, cell free supernatant.
