## Supplementary material for "Human milk oligosaccharide metabolism by *Clostridium* species suppresses inflammation and pathogen growth": fig S11

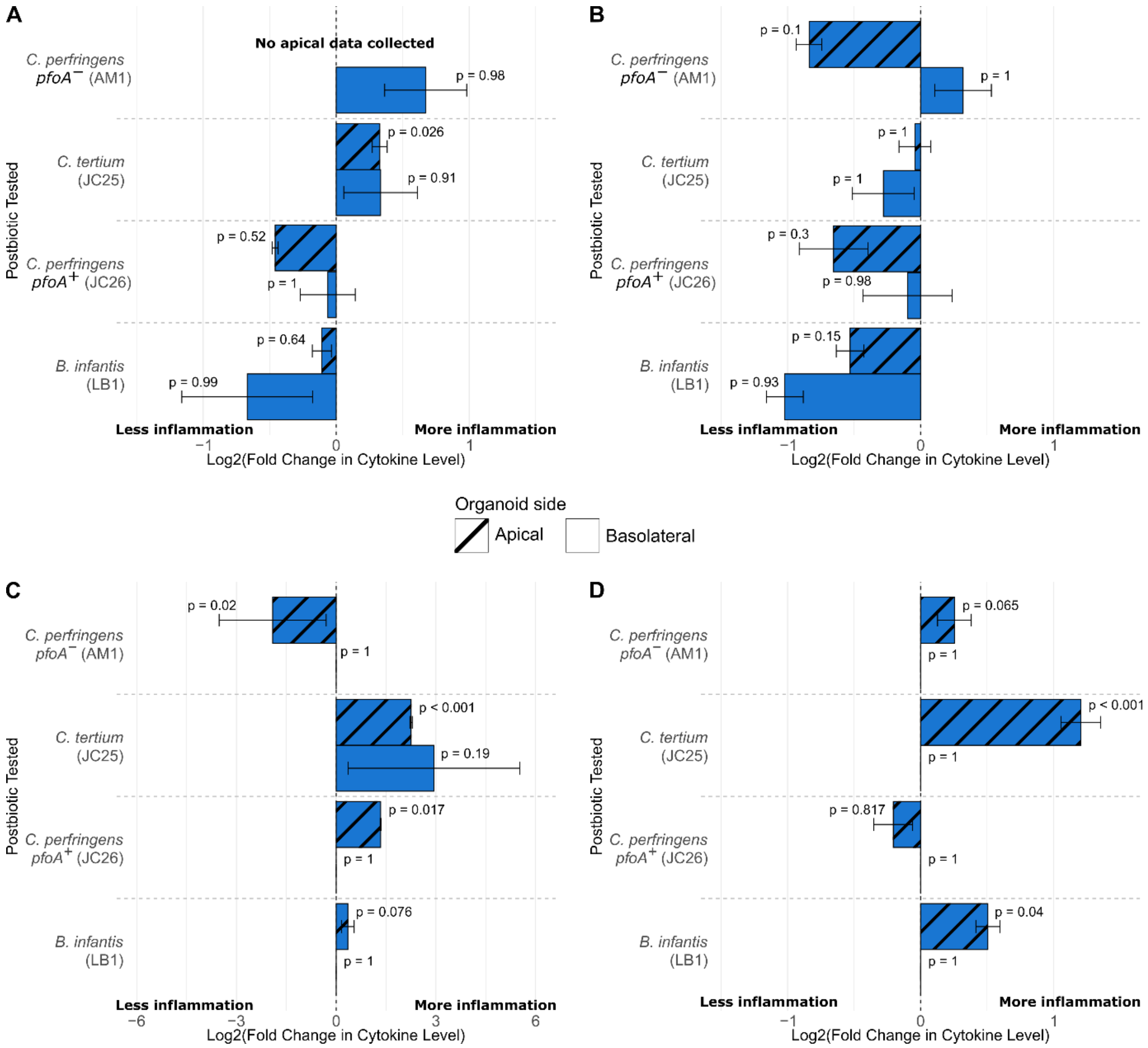


**Figure S11. Changes in organoid monolayer apical and basolateral cytokine secretion during treatment with cell free supernatants only**. Data has been converted to fold change vs the 'no treatment' control and log2 transformed for plotting. P values are for the differences between the unprocessed detected cytokine levels. (**A**) CXCL5 (**B**) CCL2 (**C**) TNF-α (**D**) CCL7.
