## Supplementary material for "Human milk oligosaccharide metabolism by *Clostridium* species suppresses inflammation and pathogen growth": table S1

**Table S1. Antibiotics resistance of selected isolates.** *Denotes resistance to the antibiotic.

| Isolate | Species | Vancomycin | Ampicillin | Metronidazole | Penicillin | Meropenem |
|---|---|---|---|---|---|---|
| AM76 | *B. breve* | ≤0.125 | 0.0625 | 1 | 0.032 | ≤0.004 |
| BM6-3 | *B. breve* | 1 | 0.5 | **>8 *** | 0.25 | >0.5 |
| CS27 | *B. bifidum* | 2 | 0.032 | 4 | 0.032 | 0.016 |
| LB1 | *B. infantis* | 1 | 0.25 | **>8 *** | 0.125 | 0.25 |
| PP1 | *B. animalis* | 0.5 | 0.032 | 2 | 0.0625 | 0.032 |
| VK68 | *B. breve* | 0.5 | 0.0625 | 1 | 0.032 | ≤0.004 |
| AM1 | *C. perfringens* | 1 | 0.016 | 2 | 0.0625 | 0.016 |
| AM37 | *C. baratii* | 2 | 0.25 | 0.5 | 0.25 | 0.032 |
| AM226 | *C. perfringens* | 1 | 0.0625 | 0.5 | 0.0625 | 0.016 |
| AM248 | *C. perfringens* | 0.5 | 0.0625 | 1 | 0.0625 | 0.016 |
| JC25 | *C. tertium* | 2 | 1 | 1 | **>1 *** | **0.5 *** |
| JC26 | *C. perfringens* | 0.5 | 0.125 | 0.5 | 0.125 | 0.016 |
| JC29 | *C. perfringens* | 0.5 | 0.125 | 1 | 0.0625 | 0.032 |
| JC31 | *C. butyricum* | 1 | 0.125 | 0.5 | 0.25 | 0.125 |
| JC33 | *C. perfringens* | 0.5 | 0.0625 | 1 | 0.125 | 0.016 |
| JC36 | *C. perfringens* | 0.5 | 0.0625 | 1 | 0.0625 | 0.008 |
| JC53 | *C. paraputrificum* | 0.5 | 0.0625 | 0.5 | 0.0625 | 0.016 |
| JC66 | *C. paraputrificum* | 1 | 0.25 | 2 | 0.0625 | 0.032 |
| TB43 | *C. tertium* | 2 | 1 | 1 | **1 *** | **0.25 *** |
| ATCC 13124 | *C. perfringens* | 1 | 0.0625 | 1 | 0.0625 | 0.016 |
| NCTC 9343 | *B. fragilis* | >1 | >4 | 0.5 | >1 | 0.125 |
| NCTC 12973 | *S. aureus* | 1 | 1 | >8 | 1 | 0.125 |
| NCTC 12241 | *E. coli* | >1 | 4 | >8 | >1 | 0.032 |
