## Supplementary material for "Human milk oligosaccharide metabolism by *Clostridium* species suppresses inflammation and pathogen growth": table S2

**Table S2. Number of differentially expressed genes (DEGs) identified in *C. perfringens* pfoA-(AM1) during growth on individual human milk oligosaccharides (HMOs) compared with growth on lactose.**

| HMO | Number of DEGs | Number upregulated | Number downregulated |
|---|---|---|---|
| DSLNT | 326 | 238 | 88 |
| LNnT | 240 | 146 | 94 |
| 6’SL | 53 | 31 | 22 |

DSLNT, Disialyllacto-N-tetraose; LNnT, Lacto-N-neotetraose; 6’SL, 6'-Sialyllactose
