## Supplementary material for "Human milk oligosaccharide metabolism by *Clostridium* species suppresses inflammation and pathogen growth": table S3

**Table S3. Degradation byproducts following growth of *C. perfringens* pfoA$^{-}$ (AM1) on lactose, Disialyllacto-N-tetraose, and 6'-Sialyllactose**

| HMO used for growth | Concentration (g/l) | | | | | | | |
|---|---|---|---|---|---|---|---|---|
| | **Lactose** | **6'SL** | **LNT** | **LNnT** | **DSLNT** | **Fucose** | **LNB** | **Sialic acid** |
| Lactose | 0.03 | n.d. | n.d. | n.d. | n.d. | n.d. | n.d. | n.d. |
| Lactose | 0.02 | n.d. | n.d. | n.d. | n.d. | n.d. | n.d. | n.d. |
| Lactose | 0.02 | n.d. | n.d. | n.d. | n.d. | n.d. | n.d. | n.d. |
| DSLNT | 0.02 | n.d. | 0.89 | n.d. | n.d. | n.d. | 0.136 | 0.201 |
| DSLNT | 0.02 | n.d. | 0.97 | n.d. | n.d. | n.d. | 0.143 | 0.239 |
| DSLNT | 0.02 | n.d. | 0.95 | n.d. | n.d. | n.d. | 0.153 | 0.275 |
| 6'SL | 0.02 | n.d. | n.d. | n.d. | n.d. | n.d. | n.d. | n.d. |
| 6'SL | 0.02 | n.d. | n.d. | n.d. | n.d. | n.d. | n.d. | 0.02 |
| 6'SL | 0.02 | n.d. | n.d. | n.d. | n.d. | n.d. | n.d. | 0.022 |

n.d., not detected; 6’SL, 6'-Sialyllactose, LNT, Lacto-N-tetraose; LNnT, Lacto-N-neotetraose; DSLNT, Disialyllacto-N-tetraose; LNB, Lacto-N-biose I
