## Supplementary material for "Human milk oligosaccharide metabolism by *Clostridium* species suppresses inflammation and pathogen growth": table S4

**Table S4. Results of per strain PERMANOVA tests comparing the metabolomes of glucose- and HMO-derived cell free supernatent**

| Strain | Species | R2 | F | P |
|---|---|---|---|---|
| AM76 | *B. breve* | 0.4889 | 3.8265 | 0.1 |
| LB1 | *B. infantis* | 0.6921 | 8.9925 | 0.1 |
| AM7 | *B. longum* | 0.3361 | 2.0246 | 0.3 |
| AM37 | *C. baratii* | 0.7465 | 11.7808 | 0.1 |
| JC53 | *C. paraputrificum* | 0.3803 | 2.4545 | 0.3 |
| AM1 | *C. perfringens* | 0.5051 | 4.0826 | 0.1 |
| JC36 | *C. perfringens* | 0.6835 | 8.6380 | 0.1 |
| JC26 | *C. perfringens* | 0.4443 | 3.1981 | 0.1 |
| JC25 | *C. tertium* | 0.6218 | 6.5773 | 0.1 |
