## Supplementary material for "Human milk oligosaccharide metabolism by *Clostridium* species suppresses inflammation and pathogen growth": table S5

**Table S5. Cytokine production (pg) for IL8 ELISA and multiplex assays performed on apical and basolateral supernatants harvested from organoid monolayers treated with *C. perfringens* pfoA- (AM1) cell free supernatant in the presence and absence of inflammatory stimuli.**

| Condition | Replicate | Side | IL8 | CCL2 | CCL7 | TNF-α | CXCL5 | MIF | IL18 | IL1RA |
|---|---|---|---|---|---|---|---|---|---|---|
| No treatment | R1 | Apical | NA | 14.10 | 0.80 | 0.08 | NA | 4313.08 | 0.29 | 33.97 |
| No treatment | R2 | Apical | 37.75 | NA | 0.69 | 0.10 | NA | 4317.98 | 0.00 | 31.21 |
| No treatment | R3 | Apical | 39.32 | 14.18 | 0.83 | 0.11 | NA | 4395.28 | 0.15 | 34.52 |
| CFS only (25% v/v) | R1 | Apical | 52.65 | 7.75 | 1.00 | 0.08 | 1.04 | 4149.72 | 0.14 | 31.87 |
| CFS only (25% v/v) | R2 | Apical | 57.33 | 7.49 | 0.84 | 0.01 | 0.98 | 3909.29 | 0.11 | 31.10 |
| CFS only (25% v/v) | R3 | Apical | 36.96 | 8.52 | 0.95 | 0.00 | 1.09 | 4140.71 | 0.00 | 40.86 |
| CFS (25% v/v)+Stimuli | R1 | Apical | NA | 9.01 | NA | 0.14 | 0.79 | 4224.22 | 0.00 | 33.37 |
| CFS (25% v/v)+Stimuli | R2 | Apical | 100.59 | 8.37 | 1.09 | 0.10 | 1.04 | 4237.49 | 0.21 | 32.88 |
| CFS (25% v/v)+Stimuli | R3 | Apical | 109.03 | 11.50 | 1.11 | 0.12 | 1.78 | 4484.13 | 1.35 | 134.97 |
| Stimuli only | R1 | Apical | 189.54 | 63.13 | 1.90 | 0.17 | NA | 4084.82 | 0.11 | 30.87 |
| Stimuli only | R2 | Apical | 177.46 | 67.32 | NA | 0.16 | NA | 4080.50 | 0.27 | 37.13 |
| Stimuli only | R3 | Apical | 171.41 | 74.03 | 1.88 | 0.18 | NA | 4360.75 | 0.20 | 27.77 |
| No treatment | R1 | Basolateral | 6.33 | 0.89 | 0.00 | 0.00 | NA | 11951.99 | 0.00 | 28.51 |
| No treatment | R2 | Basolateral | 11.76 | 1.25 | 0.00 | 0.00 | 7.88 | 12493.45 | 0.00 | 29.29 |
| No treatment | R3 | Basolateral | 8.93 | 1.19 | 0.00 | 0.00 | 7.80 | 12881.81 | 0.41 | 36.30 |
| CFS only (25% v/v) | R1 | Basolateral | 17.36 | 1.61 | 0.00 | 0.00 | 15.90 | 12166.14 | 0.43 | 28.73 |
| CFS only (25% v/v) | R2 | Basolateral | 13.80 | 1.38 | 0.00 | 0.00 | 11.64 | 11363.71 | 0.00 | 29.99 |
| CFS only (25% v/v) | R3 | Basolateral | 10.99 | 1.20 | 0.00 | 0.00 | 10.56 | 11799.51 | 0.00 | 26.51 |
| CFS (25% v/v)+Stimuli | R1 | Basolateral | 659.36 | 2.55 | 0.00 | 0.00 | NA | 12522.85 | 0.00 | 49.97 |
| CFS (25% v/v)+Stimuli | R2 | Basolateral | 688.34 | 2.19 | 0.00 | 0.00 | 30.01 | 12205.55 | 0.50 | 31.82 |
| CFS (25% v/v)+Stimuli | R3 | Basolateral | 536.39 | 2.62 | 0.00 | NA | 29.66 | 12483.59 | 0.81 | 42.06 |
| Stimuli only | R1 | Basolateral | 1444.54 | NA | NA | 0.42 | 55.73 | 10227.08 | 0.19 | 34.33 |
| Stimuli only | R2 | Basolateral | 1370.35 | 219.45 | 8.66 | 0.13 | 96.84 | 11882.84 | 0.48 | 35.48 |
| Stimuli only | R3 | Basolateral | 1547.86 | 212.64 | 8.86 | 0.25 | 77.96 | 11924.56 | 0.00 | 30.86 |

‘NA’ represents values removed as outliers per modified z score calculation.
