## Supplementary material for "Human milk oligosaccharide metabolism by *Clostridium* species suppresses inflammation and pathogen growth": table S6

**Table S6. Cytokine production (pg) for IL8 ELISA and multiplex assays performed on apical and basolateral supernatants harvested from organoid monolayers treated with *C. perfringens* pfoA+ (JC26) cell free supernatant in the presence and absence of inflammatory stimuli.**

| Condition | Replicate | Side | IL8 | CCL 2 | CCL 7 | TNF-α | CXCL 5 | MIF | IL18 | IL1RA |
|---|---|---|---|---|---|---|---|---|---|---|
| No treatment | R1 | Apical | 37.10 | 29.10 | 1.82 | NA | 133.59 | 8505.91 | 0.00 | 20.83 |
| No treatment | R2 | Apical | 66.19 | 21.26 | NA | 0.07 | 89.02 | 12896.23 | 0.62 | 48.82 |
| No treatment | R3 | Apical | 47.22 | 33.51 | 1.83 | 0.08 | 125.51 | 11101.43 | 0.59 | 34.22 |
| CFS only (25% v/v) | R1 | Apical | 66.19 | 21.70 | 1.76 | 0.22 | NA | 9023.75 | 0.41 | 34.31 |
| CFS only (25% v/v) | R2 | Apical | 94.47 | 15.37 | 1.44 | 0.22 | 83.47 | 6028.50 | 0.13 | 12.47 |
| CFS only (25% v/v) | R3 | Apical | 77.76 | 16.80 | 1.55 | NA | 85.39 | 7612.45 | 0.33 | 27.50 |
| CFS (25% v/v)+Stimuli | R1 | Apical | NA | 23.25 | 1.85 | 0.29 | 107.44 | 8289.85 | 0.53 | 34.27 |
| CFS (25% v/v)+Stimuli | R2 | Apical | 68.87 | 10.31 | 0.98 | NA | 52.82 | 8228.28 | 0.24 | 30.35 |
| CFS (25% v/v)+Stimuli | R3 | Apical | 72.43 | 20.19 | 1.68 | 0.28 | 98.00 | 15001.60 | 0.89 | 80.60 |
| Stimuli only | R1 | Apical | 79.53 | 43.65 | 1.47 | 0.15 | 130.64 | 7999.80 | 0.33 | 18.85 |
| Stimuli only | R2 | Apical | NA | 38.12 | 1.76 | NA | 100.32 | 8850.59 | 0.22 | 20.03 |
| Stimuli only | R3 | Apical | 80.41 | 22.52 | 1.00 | 0.15 | 51.44 | 7619.28 | 0.11 | 15.72 |
| No treatment | R1 | Basolateral | 20.93 | 1.42 | 0.00 | 0.00 | 16.81 | 22459.04 | 0.00 | 22.06 |
| No treatment | R2 | Basolateral | 15.35 | NA | 0.00 | 0.00 | NA | 18888.06 | 0.00 | 23.44 |
| No treatment | R3 | Basolateral | 34.10 | 1.55 | 0.00 | 0.00 | 16.93 | 20226.99 | 0.00 | 18.66 |
| CFS only (25% v/v) | R1 | Basolateral | 23.54 | 1.49 | 0.00 | 0.00 | 15.51 | 20186.80 | 0.00 | 8.91 |
| CFS only (25% v/v) | R2 | Basolateral | 16.24 | 1.06 | 0.00 | 0.00 | 14.34 | 19762.30 | 0.00 | NA |
| CFS only (25% v/v) | R3 | Basolateral | 14.16 | 1.67 | 0.00 | 0.00 | 18.93 | 19472.58 | 0.13 | 15.21 |
| CFS (25% v/v)+Stimuli | R1 | Basolateral | 43.91 | 1.46 | 0.00 | 0.00 | 20.30 | 20639.74 | 0.24 | 21.44 |
| CFS (25% v/v)+Stimuli | R2 | Basolateral | 110.85 | 1.37 | 0.00 | 0.00 | 30.75 | 19469.69 | 0.00 | 9.61 |
| CFS (25% v/v)+Stimuli | R3 | Basolateral | 69.09 | 1.41 | 0.00 | 0.00 | 23.03 | 20684.17 | 0.29 | 25.91 |
| Stimuli only | R1 | Basolateral | 163.94 | NA | 2.12 | 0.14 | 105.39 | 21168.10 | 0.25 | 20.64 |
| Stimuli only | R2 | Basolateral | 170.08 | 29.34 | 1.99 | 0.10 | 92.95 | 20884.96 | 0.00 | 25.98 |
| Stimuli only | R3 | Basolateral | NA | 28.99 | 1.94 | 0.17 | 96.60 | 21586.98 | 0.00 | 16.81 |

‘NA’ represents values removed as outliers per z score calculation.
