## Supplementary material for "Human milk oligosaccharide metabolism by *Clostridium* species suppresses inflammation and pathogen growth": table S7

**Table S7. Cytokine production (pg) for IL8 ELISA and multiplex assays performed on apical and basolateral cell free supernatants harvested from organoid monolayers exposed to *Clostridium tertium* (JC25) CFS.**

| Condition | Replicate | Side | IL8 | CCL2 | CCL7 | TNF-α | CXCL5 | MIF | IL18 | IL1RA |
|---|---|---|---|---|---|---|---|---|---|---|
| No treatment | R1 | Apical | 61.87 | NA | 1.00 | 0.12 | NA | 9253.57 | 0.00 | 9.62 |
| No treatment | R2 | Apical | 41.09 | 25.36 | 1.23 | 0.15 | 144.25 | 11498.48 | 0.18 | 15.92 |
| No treatment | R3 | Apical | 47.98 | 25.02 | 1.34 | 0.17 | 147.42 | 11158.86 | 0.29 | 20.74 |
| CFS only (25% v/v) | R1 | Apical | 266.83 | 26.19 | 2.87 | 0.71 | 190.01 | 14932.16 | 0.86 | 40.31 |
| CFS only (25% v/v) | R2 | Apical | 238.21 | 25.06 | 2.99 | 0.73 | 175.88 | 10034.26 | 0.37 | 21.43 |
| CFS only (25% v/v) | R3 | Apical | 216.64 | 22.30 | 2.46 | 0.74 | 183.53 | 8912.46 | 0.36 | 17.79 |
| CFS (25% v/v)+Stimuli | R1 | Apical | 228.57 | 31.30 | 2.92 | 0.87 | 175.04 | 9482.99 | 0.36 | 19.25 |
| CFS (25% v/v)+Stimuli | R2 | Apical | NA | 30.84 | 3.02 | 0.69 | NA | 15418.75 | 0.98 | 51.20 |
| CFS (25% v/v)+Stimuli | R3 | Apical | 226.26 | NA | 2.53 | 0.52 | 179.22 | 8067.47 | 0.26 | 10.30 |
| Stimuli only | R1 | Apical | 100.42 | 40.69 | 1.71 | 0.09 | 161.41 | 9462.41 | 0.26 | 13.48 |
| Stimuli only | R2 | Apical | 94.08 | 44.63 | 1.81 | 0.10 | 146.53 | 10375.10 | 0.00 | 12.62 |
| Stimuli only | R3 | Apical | 90.35 | 38.82 | 1.97 | NA | 143.44 | 9163.73 | 0.07 | 11.54 |
| No treatment | R1 | Basolateral | 15.71 | 1.99 | 0.00 | 0.00 | 20.86 | 28464.35 | 0.00 | 10.24 |
| No treatment | R2 | Basolateral | 17.49 | 1.93 | 0.00 | 0.00 | 19.79 | NA | 0.00 | 9.41 |
| No treatment | R3 | Basolateral | 23.34 | NA | NA | NA | 19.32 | 24383.91 | 0.00 | 8.80 |
| CFS only (25% v/v) | R1 | Basolateral | 73.91 | 1.57 | 0.00 | 0.27 | 26.32 | 25992.97 | 0.00 | 14.33 |
| CFS only (25% v/v) | R2 | Basolateral | 76.79 | 1.91 | 0.00 | 0.00 | 29.79 | 26042.30 | 0.00 | 5.84 |
| CFS only (25% v/v) | R3 | Basolateral | NA | 1.39 | 0.00 | 0.15 | 20.42 | 24086.06 | 0.00 | 8.76 |
| CFS (25% v/v)+Stimuli | R1 | Basolateral | 135.39 | NA | 0.00 | 0.20 | 47.57 | 24888.63 | 0.00 | 10.14 |
| CFS (25% v/v)+Stimuli | R2 | Basolateral | 188.61 | 2.65 | 0.00 | NA | 59.40 | 24683.76 | 0.00 | 9.16 |
| CFS (25% v/v)+Stimuli | R3 | Basolateral | 116.49 | 2.71 | 0.00 | 0.16 | 40.64 | 24745.89 | 0.00 | 11.01 |
| Stimuli only | R1 | Basolateral | NA | 40.94 | 2.57 | 0.03 | 186.85 | 25581.08 | 0.50 | 11.64 |
| Stimuli only | R2 | Basolateral | 253.93 | 45.03 | 3.17 | 0.00 | 167.88 | 25337.50 | 0.00 | NA |
| Stimuli only | R3 | Basolateral | 254.32 | 47.13 | 3.40 | 0.06 | 203.58 | 26462.40 | 0.00 | 12.11 |

'NA' represents values removed as outliers per z score calculation.
