## Supplementary material for "Human milk oligosaccharide metabolism by *Clostridium* species suppresses inflammation and pathogen growth": table S8

**Table S8. Cytokine production (pg) for IL8 ELISA and multiplex assays performed on apical and basolateral supernatants harvested from organoid monolayers treated with *Bifidobacterium infantis* (LB1) cell free supernatant in the presence and absence of inflammatory stimuli.**

| Condition | Replicate | Side | IL8 | CCL2 | CCL7 | TNF-α | CXCL5 | MIF | IL18 | IL1RA |
|---|---|---|---|---|---|---|---|---|---|---|
| No treatment | R1 | Apical | 117.69 | NA | 1.46 | 0.51 | 383.70 | 13510.51 | 0.41 | 65.92 |
| No treatment | R2 | Apical | 140.21 | 31.04 | 1.67 | 0.52 | 383.48 | 9531.35 | 0.41 | 44.49 |
| No treatment | R3 | Apical | 221.51 | 31.10 | 1.56 | NA | NA | 11100.42 | 0.43 | 47.00 |
| CFS only (25% v/v) | R1 | Apical | 232.86 | 21.83 | 2.33 | 0.58 | 362.83 | 11360.28 | 0.60 | 62.92 |
| CFS only (25% v/v) | R2 | Apical | 337.41 | 22.94 | 2.28 | 0.75 | 369.46 | 12328.38 | 0.81 | 76.30 |
| CFS only (25% v/v) | R3 | Apical | 205.92 | 19.90 | 2.07 | 0.66 | 336.41 | 10440.34 | 0.24 | 52.34 |
| CFS (25% v/v)+Stimuli | R1 | Apical | 129.65 | 26.39 | 2.15 | 0.44 | 352.27 | 10588.72 | 0.41 | 62.36 |
| CFS (25% v/v)+Stimuli | R2 | Apical | NA | 31.75 | 2.61 | NA | NA | 9678.06 | 0.00 | 47.60 |
| CFS (25% v/v)+Stimuli | R3 | Apical | 142.33 | 24.09 | 1.95 | 0.39 | 328.16 | 10864.58 | 0.50 | 59.83 |
| Stimuli only | R1 | Apical | 198.84 | NA | NA | 0.43 | 438.57 | 10142.59 | 0.39 | 45.72 |
| Stimuli only | R2 | Apical | 203.80 | 48.64 | 2.32 | 0.35 | 353.11 | 9755.74 | 0.36 | 38.30 |
| Stimuli only | R3 | Apical | NA | 49.16 | 2.35 | 0.46 | 423.29 | 9190.79 | 0.39 | 45.88 |
| No treatment | R1 | Basolateral | 46.03 | 2.48 | 0.00 | 0.00 | 19.26 | 9525.01 | 0.43 | 37.78 |
| No treatment | R2 | Basolateral | 56.81 | 3.39 | 0.00 | 0.00 | 33.37 | 17143.96 | 0.00 | 45.71 |
| No treatment | R3 | Basolateral | 59.56 | 3.87 | 0.00 | 0.00 | 26.81 | 14279.69 | 0.00 | 33.17 |
| CFS only (25% v/v) | R1 | Basolateral | 36.65 | 1.49 | 0.00 | 0.00 | 17.31 | 13700.44 | 0.00 | 32.85 |
| CFS only (25% v/v) | R2 | Basolateral | 33.00 | 1.71 | 0.00 | 0.00 | 22.93 | 15251.86 | 0.14 | 34.17 |
| CFS only (25% v/v) | R3 | Basolateral | 38.71 | NA | 0.00 | NA | 11.63 | 17557.91 | 2.98 | 19.55 |
| CFS (25% v/v)+Stimuli | R1 | Basolateral | 319.94 | 21.30 | 0.24 | 0.00 | 87.99 | 6439.69 | 0.00 | 35.41 |
| CFS (25% v/v)+Stimuli | R2 | Basolateral | 479.03 | 29.51 | NA | 0.00 | 153.90 | 6514.37 | 0.00 | 41.48 |
| CFS (25% v/v)+Stimuli | R3 | Basolateral | 393.13 | 26.50 | 0.37 | 0.00 | 105.01 | 6061.47 | 0.00 | 30.29 |
| Stimuli only | R1 | Basolateral | 682.98 | 54.65 | NA | 32.14 | 207.25 | 17535.39 | 0.00 | 23.65 |
| Stimuli only | R2 | Basolateral | 797.05 | 52.33 | 3.05 | 39.93 | 127.51 | 17600.39 | 0.97 | 41.59 |
| Stimuli only | R3 | Basolateral | 830.37 | 46.69 | 3.80 | 34.25 | 106.92 | 16316.34 | 0.70 | 31.88 |

‘NA’ represents values removed as outliers per z score calculation.
