## Supplementary material for "Human milk oligosaccharide metabolism by *Clostridium* species suppresses inflammation and pathogen growth": table S9

**Table S9. Cytokine production (pg) for IL8 ELISA and multiplex assays performed on apical and basolateral supernatants harvested from organoid monolayers treated with ZMB1 medium in the presence and absence of inflammatory stimuli.**

| Condition | Replicate | Side | IL8 | CCL2 | CCL7 | TNF-α | CXCL5 | MIF | IL18 | IL1RA |
|---|---|---|---|---|---|---|---|---|---|---|
| No treatment | R1 | Apical | 23.98 | 18.37 | 1.99 | 0.00 | 97.04 | 7825.83 | 0.00 | 0.00 |
| No treatment | R2 | Apical | 32.45 | 24.92 | 2.05 | 0.00 | 132.73 | 5984.90 | 0.13 | 20.35 |
| No treatment | R3 | Apical | 28.21 | 19.49 | 1.98 | NA | 112.30 | 6799.74 | 0.00 | 21.44 |
| CFS only (25% v/v) | R1 | Apical | 32.10 | NA | 1.98 | 0.05 | 124.48 | 10344.96 | 0.59 | 67.92 |
| CFS only (25% v/v) | R2 | Apical | 46.60 | 17.37 | 2.47 | 0.06 | 142.85 | 5610.08 | 0.33 | 28.83 |
| CFS only (25% v/v) | R3 | Apical | 38.46 | 17.41 | 2.35 | NA | 148.27 | 5787.93 | 0.18 | 21.34 |
| CFS (25% v/v)+Stimuli | R1 | Apical | 87.49 | 25.87 | 3.11 | NA | 159.50 | 5172.83 | 0.22 | 27.98 |
| CFS (25% v/v)+Stimuli | R2 | Apical | 73.96 | 24.66 | 2.97 | 0.00 | 153.99 | 4930.55 | 0.00 | 21.76 |
| CFS (25% v/v)+Stimuli | R3 | Apical | 54.76 | NA | 2.48 | 0.00 | NA | 5856.50 | 0.05 | 29.53 |
| Stimuli only | R1 | Apical | 71.47 | 34.00 | 2.85 | NA | 150.15 | 8252.40 | 0.29 | 30.43 |
| Stimuli only | R2 | Apical | NA | 33.96 | 2.67 | 0.13 | 119.03 | 6874.74 | 0.18 | 24.76 |
| Stimuli only | R3 | Apical | 72.18 | NA | 2.54 | 0.14 | 132.39 | 9564.37 | 0.95 | 64.05 |
| No treatment | R1 | Basolateral | 14.80 | NA | 0.00 | 0.00 | 8.57 | 12308.03 | 0.00 | 35.06 |
| No treatment | R2 | Basolateral | 8.82 | 1.56 | 0.00 | 0.00 | 11.70 | 10910.58 | 0.00 | 27.85 |
| No treatment | R3 | Basolateral | 12.50 | 1.49 | 0.00 | 0.00 | 12.52 | 10952.33 | 0.00 | 37.41 |
| CFS only (25% v/v) | R1 | Basolateral | 9.17 | NA | 0.00 | 0.00 | 13.75 | 15553.69 | 0.00 | 63.12 |
| CFS only (25% v/v) | R2 | Basolateral | 9.17 | 1.24 | 0.00 | 0.00 | 8.65 | 9926.34 | 0.00 | 16.54 |
| CFS only (25% v/v) | R3 | Basolateral | NA | 1.35 | 0.00 | 0.00 | 12.40 | 10175.43 | 0.00 | 18.59 |
| CFS (25% v/v)+Stimuli | R1 | Basolateral | 415.98 | 54.00 | 8.79 | 0.21 | 77.27 | 9633.02 | 0.00 | 22.91 |
| CFS (25% v/v)+Stimuli | R2 | Basolateral | NA | 40.53 | 7.36 | NA | 59.58 | 10177.78 | 0.00 | 24.87 |
| CFS (25% v/v)+Stimuli | R3 | Basolateral | 410.05 | 42.92 | 6.62 | 0.21 | 54.25 | 10220.53 | 0.00 | 24.19 |
| Stimuli only | R1 | Basolateral | 400.80 | NA | NA | 0.15 | 46.70 | 11265.94 | 0.00 | 0.00 |
| Stimuli only | R2 | Basolateral | 418.24 | 45.91 | 7.62 | 0.16 | 78.68 | 13498.10 | 0.37 | 52.53 |
| Stimuli only | R3 | Basolateral | 452.06 | 45.57 | 7.66 | 0.18 | 70.54 | 13109.75 | 0.00 | 39.58 |

‘NA’ represents values removed as outliers per z score calculation.
