## Supplementary material for "Human milk oligosaccharide metabolism by *Clostridium* species suppresses inflammation and pathogen growth": table S10

**Table S10. Cytokine production (pg) for IL8 ELISA and multiplex assays performed on apical and basolateral supernatants harvested from organoid monolayers treated with pH adjusted *C. perfringens* pfoA- (AM1) cell free supernatent in the presence and absence of inflammatory stimuli.**

| Condition | Replicate | Side | IL8 | CCL2 | CCL7 | TNF-α | CXCL5 | MIF | IL18 | IL1RA |
|---|---|---|---|---|---|---|---|---|---|---|
| No treatment | R1 | Apical | NA | NA | NA | 0.16 | 156.67 | 14616.21 | 0.00 | NA |
| No treatment | R2 | Apical | 36.34 | 16.98 | 1.94 | NA | 147.29 | 16116.32 | 0.82 | 44.97 |
| No treatment | R3 | Apical | 34.83 | 16.10 | 2.20 | NA | 151.91 | 12166.00 | 0.26 | 28.14 |
| Acidic CFS (25% v/v) | R1 | Apical | 63.87108 | 21.74641 | 2.412577 | 0.188526 | 47.15707 | 9050.927 | 0.038697 | 19.80196 |
| Acidic CFS (25% v/v) | R2 | Apical | 38.35432 | 15.73435 | 1.709502 | 0.109675 | 33.84564 | 7695.207 | 0 | 12.8481 |
| Acidic CFS (25% v/v)+Stimuli | R1 | Apical | 31.81 | NA | NA | 0.12 | 17.57 | 10344.13 | 0.19 | 26.73 |
| Acidic CFS (25% v/v)+Stimuli | R2 | Apical | 36.84 | 22.21 | 2.43 | 0.15 | 42.48 | 7150.71 | 0.09 | 14.54 |
| Acidic CFS (25% v/v)+Stimuli | R3 | Apical | 42.91 | 21.50 | 2.41 | NA | 49.70 | 9034.92 | 0.07 | 17.98 |
| Neutral CFS (25% v/v) | R1 | Apical | 51.05 | NA | 3.05 | NA | NA | 7749.82 | 0.00 | 12.84 |
| Neutral CFS (25% v/v) | R2 | Apical | 56.17 | 15.23 | NA | 0.09 | 61.15 | 8234.44 | 0.00 | 18.55 |
| Neutral CFS (25% v/v) | R3 | Apical | 33.82 | 15.87 | 3.03 | 0.08 | 71.53 | 7355.22 | 0.08 | 16.78 |
| Neutral CFS (25% v/v)+Stimuli | R1 | Apical | 46.97 | 21.66 | NA | 0.20 | 85.31 | 13981.39 | 0.44 | 50.20 |
| Neutral CFS (25% v/v)+Stimuli | R2 | Apical | 41.89 | 11.75 | 2.48 | 0.15 | 55.14 | 7446.27 | 0.08 | 14.12 |
| Neutral CFS (25% v/v)+Stimuli | R3 | Apical | 62.33 | 15.38 | 2.53 | 0.17 | 61.24 | 8492.82 | 0.04 | 17.40 |
| Stimuli only | R1 | Apical | 102.88 | 54.55 | NA | 0.11 | 205.27 | 12333.20 | 0.26 | 35.04 |
| Stimuli only | R2 | Apical | NA | 56.00 | 2.84 | 0.19 | 207.76 | 10884.29 | 0.22 | 19.20 |
| Stimuli only | R3 | Apical | 97.64 | NA | 2.74 | 0.17 | NA | 12858.06 | 0.35 | 30.66 |
| No treatment | R1 | Basolateral | 7.10 | 1.45 | 0.00 | 0.17 | 17.51 | 27606.25 | 0.23 | 33.98 |
| No treatment | R2 | Basolateral | 8.07 | 1.01 | 0.00 | 0.09 | 13.76 | 27134.48 | 0.00 | 34.17 |
| No treatment | R3 | Basolateral | 5.66 | 1.27 | 0.00 | 0.25 | 16.87 | 25551.81 | 0.13 | 22.24 |
| Acidic CFS (25% v/v) | R1 | Basolateral | 3.767058 | 1.676653 | 0 | 0.142578 | 18.74501 | 24624.94 | 0.199471 | 22.72643 |
| Acidic CFS (25% v/v) | R2 | Basolateral | 2.061139 | 0.937293 | 0 | 0.174563 | 16.38219 | 26546.12 | 0.07769 | 22.18571 |
| Acidic CFS (25% v/v)+Stimuli | R1 | Basolateral | NA | 2.04 | 0.00 | 0.49 | 34.05 | 27261.71 | 0.29 | 23.03 |
| Acidic CFS (25% v/v)+Stimuli | R2 | Basolateral | 53.05 | NA | 0.00 | NA | 45.24 | 26450.50 | 0.00 | 22.23 |
| Acidic CFS (25% v/v)+Stimuli | R3 | Basolateral | 55.83 | 1.98 | 0.00 | 0.33 | 29.87 | 31897.91 | 0.22 | 15.92 |
| Neutral CFS (25% v/v) | R1 | Basolateral | 4.55 | 0.99 | 0.00 | 0.24 | 14.15 | 25504.56 | 0.44 | 19.15 |
| Neutral CFS (25% v/v) | R2 | Basolateral | 7.42 | 0.80 | 0.00 | 0.21 | 16.52 | 27459.47 | 0.16 | 28.88 |
| Neutral CFS (25% v/v) | R3 | Basolateral | 2.99 | 1.08 | 0.00 | 0.22 | 18.65 | 28073.44 | 0.09 | NA |
| Neutral CFS (25% v/v)+Stimuli | R1 | Basolateral | 84.06 | 2.18 | 0.00 | 0.10 | 52.03 | 25165.28 | 0.18 | 29.01 |
| Neutral CFS (25% v/v)+Stimuli | R2 | Basolateral | 105.71 | 2.18 | 0.00 | 0.37 | 59.87 | 27205.41 | 0.21 | 24.17 |
| Neutral CFS (25% v/v)+Stimuli | R3 | Basolateral | 167.77 | NA | 0.00 | 0.29 | 70.57 | 26679.38 | 0.34 | 28.37 |
| Stimuli only | R1 | Basolateral | 145.88 | 48.37 | 5.34 | 0.28 | 121.97 | 28576.02 | 0.30 | 35.24 |

| | | | | | | | | | | |
|---|---|---|---|---|---|---|---|---|---|---|
| Stimuli only | R2 | Basolateral | 198.34 | 41.49 | 5.62 | NA | 98.45 | 25973.14 | 0.25 | 26.62 |
| Stimuli only | R3 | Basolateral | 172.12 | 46.79 | NA | 0.28 | 90.87 | 24036.04 | 0.38 | 24.59 |

‘NA’ represents values removed as outliers per z score calculation.
