## Supplementary material for "Human milk oligosaccharide metabolism by *Clostridium* species suppresses inflammation and pathogen growth": table S11

**Table S11. Cytokine production (pg) for IL8 ELISA and multiplex assays performed on apical and basolateral supernatants harvested from organoid monolayers treated with live *C. perfringens* pfoA- (AM1) and *C. perfringens* pfoA+ (JC26) in the presence and absence of inflammatory stimuli.**

| Condition | Replicate | Side | IL8 | CCL2 | CCL7 | TNF-α | CXCL5 | MIF | IL18 | IL1RA |
|---|---|---|---|---|---|---|---|---|---|---|
| AM1 only | R1 | Apical | 33.62 | 5.49 | 0.00 | 0.00 | 0.00 | 23944.36 | 1.89 | 86.48 |
| AM1 only | R2 | Apical | 27.02 | NA | 0.00 | 0.14 | 0.00 | 21610.20 | 1.12 | 76.77 |
| AM1 only | R3 | Apical | 47.68 | 5.51 | 0.00 | 0.12 | 0.00 | 27301.64 | 4.22 | 117.76 |
| AM1 + JC26 | R1 | Apical | 29.50 | 4.06 | 0.00 | 0.11 | 0.00 | 49097.00 | 8.94 | 233.25 |
| AM1 + JC26 | R2 | Apical | 27.02 | NA | 0.00 | 0.00 | 0.00 | 45046.09 | 5.18 | 175.70 |
| AM1 + JC26 | R3 | Apical | 18.79 | 4.14 | 0.00 | 0.18 | 0.00 | NA | NA | NA |
| AM1 + Stimuli | R1 | Apical | 44.37 | 6.09 | 0.00 | 0.05 | 0.00 | 23089.56 | 1.43 | NA |
| AM1 + Stimuli | R2 | Apical | 36.10 | 6.04 | 0.00 | 0.12 | 0.00 | 20479.69 | 2.85 | 104.20 |
| AM1 + Stimuli | R3 | Apical | 46.02 | NA | 0.00 | 0.20 | 0.00 | 22458.88 | 3.31 | 102.81 |
| JC26 only | R1 | Apical | 22.90 | 4.35 | 0.00 | 0.05 | 0.07 | 98212.41 | 22.63 | 465.16 |
| JC26 only | R2 | Apical | NA | 4.43 | 0.00 | 0.00 | 0.17 | 97453.40 | 25.38 | 478.98 |
| JC26 only | R3 | Apical | 23.72 | NA | 0.00 | 0.03 | NA | NA | 23.59 | 436.53 |
| No treatment | R1 | Apical | 27.02 | 12.74 | 1.01 | 0.09 | NA | 18099.16 | 0.49 | 74.66 |
| No treatment | R2 | Apical | 25.37 | 9.50 | 0.76 | NA | 116.45 | 16954.49 | 0.53 | 51.97 |
| No treatment | R3 | Apical | 34.45 | 10.48 | 0.89 | 0.09 | 111.73 | 20190.59 | NA | 82.92 |
| Stimuli only | R1 | Apical | 40.23 | 9.59 | 0.82 | 0.08 | 103.31 | NA | 0.78 | 54.23 |
| Stimuli only | R2 | Apical | 41.88 | 10.28 | 0.90 | 0.10 | 91.85 | 14465.95 | 0.65 | 52.75 |
| Stimuli only | R3 | Apical | 37.75 | 9.80 | 0.87 | 0.10 | 94.16 | 18785.79 | 0.87 | 61.21 |
| AM1 only | R1 | Basolateral | 29.24 | 1.39 | 0.00 | 0.00 | 35.72 | 80225.84 | 4.87 | 303.00 |
| AM1 only | R2 | Basolateral | 22.77 | 1.47 | 0.00 | 0.00 | 32.69 | 74186.62 | 5.19 | 254.92 |
| AM1 only | R3 | Basolateral | 30.60 | NA | 0.00 | 0.00 | 19.61 | 72761.13 | 5.77 | 283.14 |
| AM1 + JC26 | R1 | Basolateral | NA | NA | 0.00 | 0.00 | 21.64 | 68866.96 | 4.29 | NA |
| AM1 + JC26 | R2 | Basolateral | 29.79 | 1.54 | 0.00 | 0.00 | NA | 71599.08 | 5.49 | 334.37 |
| AM1 + JC26 | R3 | Basolateral | 31.95 | 1.48 | 0.00 | 0.00 | 20.12 | NA | 9.21 | 538.41 |
| AM1 + Stimuli | R1 | Basolateral | 24.12 | 0.82 | 0.00 | 0.00 | 24.75 | 67646.34 | 4.27 | 275.65 |
| AM1 + Stimuli | R2 | Basolateral | 22.50 | NA | 0.00 | 0.00 | 22.58 | NA | 3.96 | 294.97 |
| AM1 + Stimuli | R3 | Basolateral | 19.53 | 0.81 | 0.00 | 0.00 | 21.55 | 66583.61 | 3.43 | 257.11 |
| JC26 only | R1 | Basolateral | 17.92 | 1.31 | 0.00 | 0.00 | 28.43 | 72237.55 | 5.90 | 302.06 |
| JC26 only | R2 | Basolateral | 6.37 | 0.93 | 0.00 | 0.00 | 13.98 | 69695.54 | 5.98 | 273.01 |
| JC26 only | R3 | Basolateral | 12.81 | 1.18 | 0.00 | 0.00 | 26.08 | 80773.05 | NA | 290.85 |
| No treatment | R1 | Basolateral | 22.23 | 0.97 | 0.00 | 0.00 | 29.92 | 64605.95 | 2.87 | 201.39 |
| No treatment | R2 | Basolateral | 26.81 | 1.15 | 0.00 | 0.00 | 33.86 | 71296.08 | 3.87 | 274.90 |
| No treatment | R3 | Basolateral | 32.49 | 1.21 | 0.00 | 0.00 | 35.82 | 87782.00 | 5.44 | 360.67 |
| Stimuli only | R1 | Basolateral | 38.71 | 1.19 | 0.00 | 0.00 | NA | 73786.31 | 4.83 | 309.98 |
| Stimuli only | R2 | Basolateral | NA | 0.65 | 0.00 | 0.00 | 24.15 | 75376.21 | 4.55 | 284.99 |
| Stimuli only | R3 | Basolateral | 39.52 | 0.86 | 0.00 | 0.00 | 24.06 | 71717.99 | NA | 319.18 |

‘NA’ represents values removed as outliers per z score calculation.
